# Prolonged Pain Reliably Slows Peak Alpha Frequency by Reducing Fast Alpha Power

**DOI:** 10.1101/2021.07.22.453260

**Authors:** Andrew J. Furman, Mariya Prokhorenko, Michael L. Keaser, Jing Zhang, Shuo Chen, Ali Mazaheri, David A. Seminowicz

## Abstract

The relationship between the 8-12 Hz “alpha: rhythm, the predominant oscillatory activity of the brain, and pain remains unclear. In healthy individuals, acute, noxious stimuli suppress alpha power while patients with chronic pain demonstrate both enhanced alpha power and slowing of the peak alpha frequency (PAF). To investigate these apparent differences, EEG was recorded from healthy individuals while they completed two models of prolonged pain, Phasic Heat Pain and Capsaicin Heat Pain, at two testing visits occurring roughly 8 weeks apart. We report that PAF is reliably slowed and that alpha power is reliably decreased in response to prolonged pain. Furthermore, we show that alpha power changes, but not PAF changes, are fully reversed with stimulus removal suggesting that PAF slowing reflects pain associated states such as sensitization rather than the presence of ongoing pain. Finally, we provide evidence that changes to alpha power and PAF are due to power decreases in the “fast” (10-12 Hz) range of the alpha rhythm. This frequency dependent pain response aligns with the hypothesis that the alpha rhythm is composed of multiple, independent oscillators, and suggest that modulation of a putative “fast” oscillator may represent a promising therapeutic target for treating ongoing pain. In sum, we provide strong evidence that PAF is reliably slowed during prolonged pain and additionally identify a mechanism, “fast” alpha Power, which is responsible for these PAF changes.

---

The 8-12 Hz “alpha” rhythm is the predominant oscillatory activity observed over human and primate sensory cortices and is thought to reflect processes associated with the phasic inhibition of mesoscopic, neuronal activity (e.g. Haegens et al., 2011). In the spectral domain, alpha activity can be roughly described by two key features: 1) the specific frequency at which maximum power is expressed, the so called peak or individual alpha frequency (PAF/IAF), and 2) the amount of power present across the entire alpha range (alpha power).

Given its putative role in sensory processing (Ergenoglu et al., 2004; Klimesch et al., 2006; Jensen & Mazaheri, 2010; Foxe & Snyder, 2011), the alpha rhythm has received considerable attention in studies of pain and nociception (e.g. Babiloni et al., 2006). Most studies have focused on alpha power and reported that experimental models of phasic pain are associated with the suppression of alpha power (Dowman et al., 2008, Peng et al., 2014, Huishi Zhang et al., 2016). On the other hand, lower frequency range alpha power (<10 Hz) is enhanced in patients with chronic pain suggesting that alpha power may be a useful diagnostic for separating acute and prolonged pain (Sarnthein et al., 2006).

More recently, attention has turned to PAF as abnormalities have been observed in a wide variety of chronic pain conditions (Sarnthein et al., 2006, Walton et al., 2010, de Vries et al., 2013, Kim et al., 2019). Despite a lack of evidence that PAF slowing follows chronic pain emergence, PAF slowness has been interpreted to reflect disease processes involved in the transition to and persistence of chronic pain (Walton & Llinás, 2010). An alternative explanation, however, is that slow PAF is a pre-injury neural feature that reflects heightened pain sensitivity and subsequently predisposes an individual to developing chronic pain (Furman et al., 2018). Indeed, work from our lab has shown that PAF collected prior to a painful event can predict the future amount of pain a healthy individual will experience (i.e. their pain sensitivity). This relationship exists across several different pain modalities and paradigms and is reliable across multiple timepoints (Furman et al., 2018, 2019, 2020). Clarifying these two alternatives can determine whether slow PAF is better served as a prospective diagnostic of risk or a metric of ongoing chronic pain.

To investigate how prolonged pain impacts alpha oscillations, we collected EEG data during two experimental models of prolonged pain occurring at two procedurally identical visits separated by 8 weeks on average. We aimed to determine: 1) whether PAF is slowed in response to prolonged pain; 2) whether alpha power is increased in response to prolonged pain; and 3) the relationship between PAF and alpha power changes during prolonged pain. With respect to this last objective, resolving how alpha power changes produce PAF changes may reveal novel alpha mechanisms that can be leveraged therapeutically to either treat pain or prevent the transition from acute to chronic pain.

## Materials and methods

### Participants

Sixty-one pain-free, adult participants (31 males, mean age = 27.82, age range = 21-42) without history of neurological or psychiatric disorder took part in the experiment between 7/6/2016 and 10/20/2017. This study was approved by the University of Maryland, Baltimore Institutional Review Board, and informed written consent was obtained from each participant prior to any study procedures. The study was pre-registered on ClinicalTrials.gov (NCT02796625), and the primary and secondary outcomes were previously reported, demonstrating a reliable relationship between pain-free PAF and prolonged pain sensitivity (Furman et al., 2020).

Table 1 provides information regarding how many participants contributed data to each analysis. For a few participants, we were unable to collect data for a particular EEG session or a technical malfunction prevented us from accurately matching their EEG and pain rating data.

**Table 1.** Overview of available datasets for each EEG session. Numbers reflect relevant datasets for Visit 1 / Visit 2.

|  | **Pain-Free 1** | **PHP** | **Pain-Free 2** | **Incubation** | **CHP** | **Pain-Free 3** | **CHP Rekindle** |
| --- | --- | --- | --- | --- | --- | --- | --- |
| *Excluded* | 4 / 3 | 4 / 3 | 4 / 3 | 4 / 3 | 4 / 3 | 4 / 3 | 4 / 3 |
| *Missing* | 0 / 0 | 4 / 3 | 1 / 0 | 7 / 4 | 1 / 0 | 6 / 1 | 7 / 4 |
| *Available* | 57 / 43 | 53 / 40 | 56 / 43 | 50 / 39 | 56 / 43 | 51 / 42 | 50 / 39 |

### EEG

Scalp EEG was collected from 63 channel BrainVision actiCAP system (Brain Products GmbH, Munich, Germany) labeled according to an extended international 10–20 system (Oostenveld and Praamstra, 2001). All electrodes were referenced online to the average across all recording channels and a common ground set at the AFz site. Electrode impendences were maintained below 5 kΩ throughout the experiment. Brain activity was continuously recorded within a 0.01–100 Hz bandpass filter, and with a digital sampling rate of 500 Hz. The EEG signal was amplified and digitized using an actiCHamp DC amplifier (Brain Products GmbH, Munich, Germany) linked to BrainVision Recorder software (version 2.1, Brain Products GmbH, Munich, Germany).

### Thermal Stimulator

Thermal stimuli were delivered to the volar surface of the participant’s left forearm using a thermal contact heat stimulator (27mm diameter Medoc Pathway CHEPS Peltier device; Medoc Advanced Medical Systems Ltd., Ramat Yishai, Israel).

### Quantitative Sensory Testing

Participants completed four threshold tests, in which they were asked to report when: (1) they felt a temperature increase (Warmth Detection Threshold); (2) they felt a temperature decrease (Cool Detection Threshold); (3) an increasing temperature first became painful (Heat Pain Threshold); and (4) a decreasing temperature first became painful (Cold Pain Threshold). A total of three trials were presented for each test with an ISI of 4-6 seconds (randomly determined on a per trial basis). Participants provided feedback for each test by clicking a computer mouse button placed in their right hand. For each test, temperatures were applied with a rise rate of 1°C/second and return rate of 2°C/second.

All sensory testing was performed on the volar surface of the left forearm. The distance from the wrist to elbow joint was measured and the forearm was divided into three equal length zones. For each test, the first trial was administered to the zone closest to the wrist, the second trial administered to the middle forearm zone, and the third trial administered to the zone closest to the elbow.

### Phasic Heat Pain (PHP) Model

Temperatures used during the PHP model were determined during each participant’s initial screening visit to the laboratory (Visit 0). During these sessions, participants were exposed to 12, 20 second trials in which a single temperature (2.5 second rise and fall) was applied to the volar surface of the left forearm. At the conclusion of each trial, participants reported the average pain they experienced during temperature application; participants were instructed to report pain ratings on a scale of 0-10, with 0 indicating no pain and 10 the most pain imaginable. Temperatures of 37, 40, 42, 44, 46 and 48°C and were each presented twice in a pseudorandom order. Trials were separated by 10 seconds and after each trial the thermode was moved to a neighboring forearm zone in order to minimize sensitization. Using pain reports from these trials, the temperature that most closely evoked an average pain rating of 5/10 was selected. This level of pain was targeted in order to best match the intensity of pain evoked by the CHP model (Furman et al., 2018). For a few participants, none of the applied temperatures could evoke a pain rating that reached 5/10. For these individuals, 48°C was used during PHP testing.

The PHP model consisted of a series of five consecutive stimulus trains. Each train lasted one minute and consisted of application of a predetermined temperature for 40 seconds (rise and fall times of 2s) followed by application of a neutral skin temperature stimulus (32°C) for 20 seconds. PHP scores were calculated by averaging pain ratings from the five, forty second periods in which the temperature was present (including temperature rise and return).

### Capsaicin Heat Pain (CHP) Model

The CHP model lasts for hours to days and recapitulates some cardinal sensory aspects of chronic neuropathic pain (Culp et al., 1989; LaMotte RH, et al.,1992; Baron 2009; Lötsch et al., 2015) without causing lasting tissue damage (Henriques & Moritz, 1947). CHP procedures were similar to those used in our prior study (Furman et al., 2018). In brief, we applied ∼1 g 10% capsaicin paste (Professional Arts Pharmacy, Baltimore, MD) topically to the volar surface of the left forearm, fixing it in place with a Tegaderm bandage. A thermode was then placed on top of the capsaicin application, heated to 40°C and held in place for 20 minutes to allow for capsaicin incubation. Given that pain from topically applied capsaicin varies as a function of skin temperature (Anderson et al., 2002), the thermode temperature was held at 40°C for all participants. This temperature was selected because, in the absence of capsaicin, all participants had heat pain thresholds higher than this temperature. CHP scores were calculated by averaging ratings across the entire five-minute CHP test that followed incubation.

To assess how reliable CHP-induced changes to PAF and alpha power, we included a “rekindling” phase (CHP Rekindle; Dirks et al., 2003). After the initial CHP testing was completed, an icepack (see below for details) was applied to the forearm until a complete termination of pain was reported. Afterwards, the thermode was again placed over top of the site of capsaicin application, heated to 40°C, and held in place for five minutes. CHP rekindle scores were calculated as the average of the pain ratings provided during this five-minute period.

### Icepack Application

At the conclusion of the PHP and CHP tests, the thermode was removed and a disposable icepack was applied the stimulated area of the left forearm. This was done to prevent pain carryover from one test to another and to ensure that pain ratings for subsequent tests were captured from a starting state of no ongoing pain. The icepack was left in place until the complete absence of pain was reported by the participant. No participants indicated that the icepack itself was ever painful. Following each icepack application and removal, a 5-minute pain-free, eyes closed EEG session occurred.

### Pain Ratings

Pain ratings were collected continuously with a manual analog scale consisting of a physical sliding tab (Medoc Advanced Medical Systems Ltd., Ramat Yishai, Israel). Prior to testing, participants were instructed that the lower and upper bounds of the scale represented no pain and the most pain imaginable, respectively, and that they should continuously update the slider to indicate the amount of pain currently being experienced. Care was taken by experimenters to avoid providing numerical anchors when describing the scale and no additional physical landmarks were present on the scale. Prior studies have found that analog scales are superior to numerical scales for capturing the pain power function often encountered in psychophysical testing (Price et al., 1994; Nielsen et al., 2005). Prior to testing, participants were given an opportunity to practice using the device with their eyes open and closed. During testing, participants were permitted to briefly open their eyes while rating. Pain ratings were collected from the manual analog scale at a rate of 1000 Hz. Manual analog scale data was transformed by converting the horizontal position of the slider into a continuous value between 0 and 100.

### Procedure

An outline of the experimental timeline and procedures is presented in Figure 1. In order to allow sufficient time for any long-term effects of capsaicin exposure to subside, visits were separated by 21 days or more (except for one case where a subject returned at 19 days because of a scheduling conflict; mean separation of Visit 1 and Visit 2 = 54.74 days, S.D. = 55.92 days, range = 19 – 310 days).

**Figure 1.**
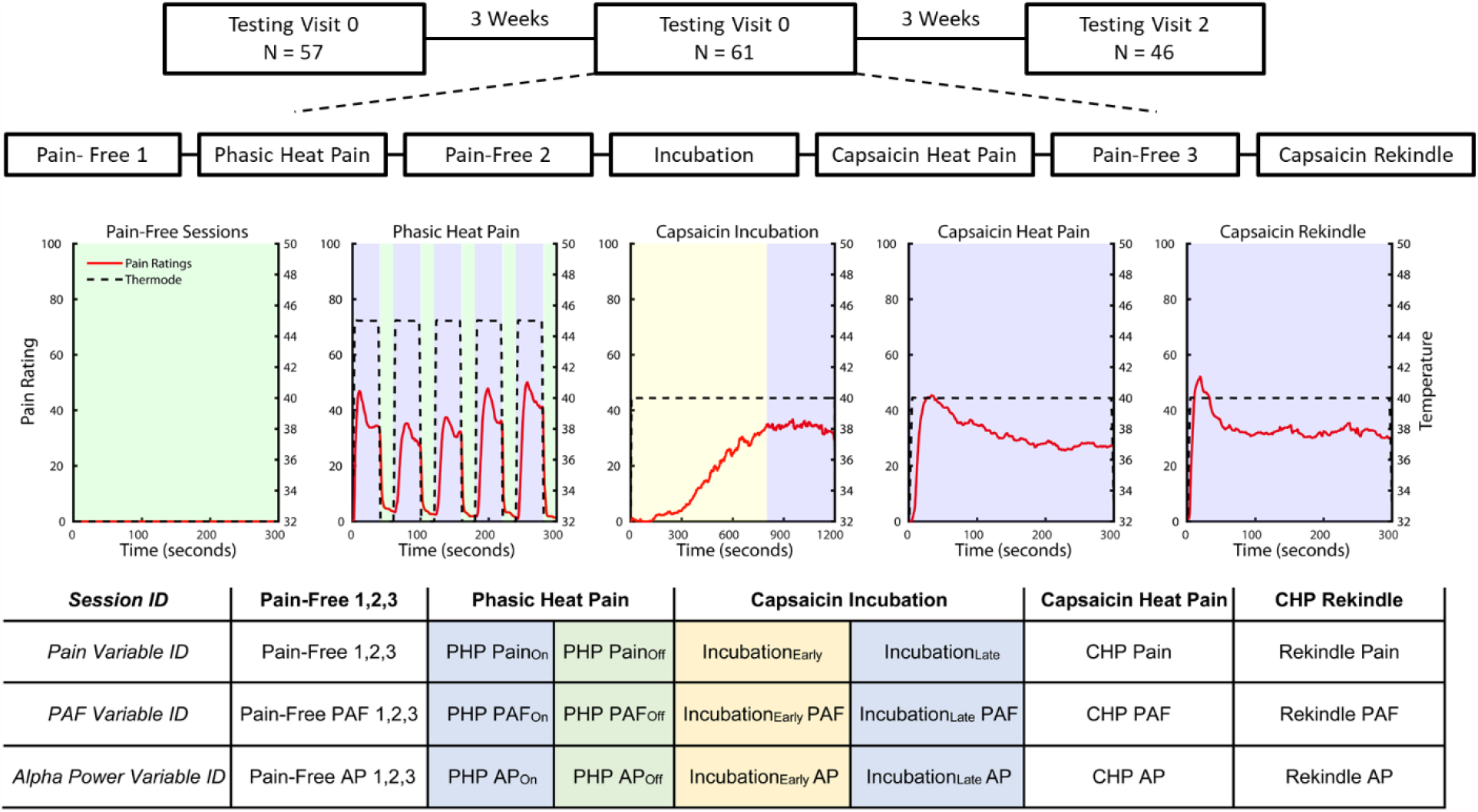
Overview of the experimental paradigm and naming conventions throughout the paper. At Testing Visits 1 and 2, participants completed the progression of experimental sessions. Prior to Pain-Free 2 & 3, an icepack was applied to the stimulated forearm for three minutes (not shown in figure). Representative data reflects the average Visit 1 pain ratings from participants who responded to capsaicin (average CHP pain > 10). Instances where EEG sessions were separated into distinct datasets are denoted by the presence of different colors within the same panel. Naming conventions for all datasets are provided in the table and, where an EEG session was divided into multiple datasets, shading color in the table has been matched to the color provided in the data panels.

Participants first underwent an initial screening visit, Visit 0, that included quantitative sensory testing as well as additional tests to ensure that 40°C was rated as minimally-painful, to identify the appropriate PHP temperature, and to provide initial exposure to capsaicin. For the first four participants, these procedures, excluding capsaicin exposure, were performed during Visit 1.

Participants returned for Visit 1 at least three weeks after completing Visit 0. Most participants then returned at least three weeks after Visit 1 for Visit 2. Procedures for Visits 1 and 2 were identical. For the entirety of Visits 1 and 2, participants were seated in a comfortable chair in a quiet room that was isolated from strong electrical interference. For all EEG sessions, lights in the testing room were turned off and participants were instructed to close their eyes, remain still, relax without falling asleep, and continuously rate any pain they experienced with the manual analog scale placed at their right hand.

Visits 1 and 2 began with quantitative sensory testing. For the first four participants, this sensory testing was not performed at Visit 2. After quantitative sensory testing, a brief 2-minute EEG was collected to ensure the quality of EEG recording. Next, a room temperature thermode was placed onto the left forearm while eyes closed, pain-free EEG was collected for 5 minutes. The relationship of PAF recorded this eyes-closed, pain-free session to pain sensitivity was reported in an earlier publication (Furman et al., 2020).

Following the pain-free EEG, prolonged pain was induced with the PHP model. During the five minutes of PHP, EEG was collected while participants rested with their eyes closed and continuously rated the intensity of pain. Upon completion of the PHP model, a disposable ice pack was placed onto the participant’s left forearm until they reported being completely free of pain after which 5 minutes of eyes closed EEG was collected. Next, the second model of prolonged pain, CHP, was induced. Participants were instructed to continuously rate pain intensity during this incubation period.

Following the 20-minute incubation period, and with the thermode temperature still held at 40°C, 5 minutes of eyes closed, continuous EEG was recorded while participants continuously rated the intensity of pain. An icepack was then applied to the forearm, removed once pain was reported to be completely absent, and followed by 5 minutes of eyes closed EEG was collected. Afterwards, a 40°C thermode was placed over the site of capsaicin application to induce CHP rekindling. Five minutes of eyes closed EEG was then recorded while participants continuously rated pain intensity.

### Data Processing

We analyzed data from the six sessions collected throughout the experiment: Pain-Free 1, PHP, Pain-Free 2, Capsaicin Incubation (Incubation), CHP, Pain-Free 3, and CHP Rekindle (Figure 1). The PHP session was further divided into periods when the noxious temperature was applied (PHP_on_) and when the skin temperature was applied (PHP_off_). Incubation data was also split into two segments based on the temporal profile of pain ratings. During the first ∼13 minutes of incubation pain ratings grew steadily (Incubation_Early_) whereas during the last ∼7 pain ratings were largely stable (Incubation_Late_).

EEG data were preprocessed with EEGLAB 13.6.5b (Delorme and Makeig, 2004). Unless otherwise noted, participants contributed 8 EEG datasets at each visit (see Table 1). For each data set, preprocessing began with filtering the data between .2 and 100Hz using a linear FIR filter. To account for potential volume conduction effects, the surface Laplacian was computed and applied to the data (Perrin et al., 1989). Finally, Independent Components Analysis (ICA) using the Infomax algorithm(may represent a promising therapeutic Bell & Sejnowski, 1995) was performed and components with spatial topographies and time series resembling blinks and/or saccades were removed from the data.

### Quantification of PAF

Frequency decomposition was performed using routines in FieldTrip (Oostenveld et al., 2011). Data from each EEG dataset was segmented into non-overlapping 5 second epochs and power spectral density in the .2–100 Hz range (0.2 Hz bins) was derived with the ‘ft_freqanalysis_mtmfft’ function. A Hanning taper was applied to the data prior to calculating the spectra to reduce edge artifacts (e.g. Mazaheri et al., 2014).

At every channel and for each epoch, PAF was estimated using a center of gravity (CoG) method (Klimesch et al., 1993). We defined CoG as follows:

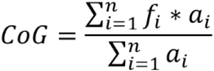

where *f*_*i*_ is the *i*th frequency bin including and above 9 Hz, n is the number of frequency bins between 9 and 11 Hz, and *a*_*i*_ the spectral amplitude for *f*_*i*_. From our previous work, we have determined that this restricted frequency range reduces the influence of 1/f EEG noise on PAF estimation (Furman et al., 2018). Epoch-level PAF estimates were averaged separately for each EEG session (i.e. Pain-free session 1, CHP session) to yield a single mean PAF estimate for each channel during each EEG session. Although our earlier work has relied exclusively on sensors putatively sampling from the sensorimotor cortices, we elected to focus our analyses on all channels given clear evidence that PAF effects are not restricted to those sensorimotor channels (Furman et al., 2018, 2019, 2020).

To ensure that results were not an artifact of the range used for PAF estimation, PAF was also calculated with the wider 8-12 Hz range. Results with this wider estimation range are provided in the supplemental section.

### Quantification of Alpha Power

Epoch-level Alpha Power (AP) estimates were calculated as the average power across the 8-12 Hz range. Epoch-level estimates from each EEG session were then averaged to yield a single AP estimate for each EEG channel during each EEG session.

## Statistical Analysis

All analyses and statistical tests were performed using custom scripts implemented in the Matlab environment (version 2019a).

For all analyses, we removed any data that was 3 standard deviations above the mean. PHP analyses were performed on all available data while CHP analyses excluded data from individuals demonstrating CHP model insensitivity. In short, we excluded participants from CHP analyses if they showed a clear pain response (average pain > 10) to PHP but not CHP.

For all analyses, linear mixed models were used to explain variance in either EEG or pain responses (see below for specifics regarding each model). To account for the large number of tests across EEG sensors, statistical differences between conditions were evaluated using non-parametric cluster-based permutation tests (Maris & Oosteveld, 2007; Pernet et al., 2015). Unless otherwise stated, tests at the single channel level were first thresholded at *p* < .05. In some cases, channels were required to surpass thresholds for multiple comparisons (i.e. a conjunction analysis) in order to move on to the cluster forming stage (see below for details specific to each analysis). Next, spatially adjacent channels surpassing this threshold were grouped into clusters based on the requirements that 1) a cluster be composed of at least two sensors and 2) that each sensor within a cluster be immediately neighbored by at least one sensor from the overall cluster.

After cluster formation, test statistic values (i.e. *t* or *F* statistics) were summed across all cluster channels to produce a single, composite test score for the entire cluster. For each cluster, this composite score was then compared against a null distribution of 1000 iteratively generated composite scores produced by randomly shuffling dependent variable values across participants and conditions at each EEG sensor belonging to the cluster in question. For *t* tests, clusters were evaluated as significant if their composite test statistic surpassed the 97.5^th^ percentile of the relevant tail of the null distribution. For *F* tests, clusters were evaluated as significant if their composite test statistic surpassed the 95^th^ percentile of the null distribution.

Where applicable, the effect size of significant fixed effects of interest (i.e. Session or PAF/Alpha Power) were calculated using Cohen’s *f*^2^ (Selya et al., 2012). We refer to results as having small, medium, or large effect sizes according to Cohen’s originally guidelines (Cohen 2013), where .02 > *f*^2^ < .15 is considered a small effect, .15 > *f*^2^ < .3 is a medium effect, and *f*^2^ > .35 is a large effect.

### Assess Whether PAF is Slowed in Response to Prolonged Pain

Seperately for PHP and CHP, PAF estimates from each channel were submitted to a linear mixed effects model with Session and Visit as fixed effects (no interaction specified) and a random effects term for Participant nested within Visit. For PHP models, clustering was performed on channels meeting the following criteria: 1) a significant (*p* < .05) difference between PHP_On_ and Pain-Free 1, and 2) a significant *a priori* contrast (*p* < .05) of PHP_On_ versus PHP_Off_ and Pain-Free 2. For CHP models, clustering was performed on channels demonstrating the following: 1) a significant difference (*p* < .05) between CHP and Pain-Free 2, and 2) a significant *a priori* contrast (*p* < .05) of CHP Rekindle versus Pain-Free 3. Cluster-based analyses were performed for both fixed effects (*t* values) and contrasts (*F* values) and only clusters surpassing the null distribution threshold for both tests were evaluated as significant.

To evaluate whether PAF is modulated by motor activity, we correlated PAF changes occurring during PHP_On_ with an estimate of slider movement. Slider movement was calculated by taking the sum of the absolute, first derivative of PHP pain ratings. Prior to this calculation, pain ratings were z-scored in order to best isolate slider movement from pain intensity.

Likelihood-ratio tests were used to assess effect stability by comparing model results against an otherwise identical model including an additional term for Session X Visit interaction. Failure to find a significant difference was taken as evidence of effect reliability across Visits.

Channels contributing to significant clusters to both PHP and CHP analyses were identified with a conjunction analysis. These results are presented for illustrative purposes and should not be used draw conclusions about any EEG channel given that significance was tested solely at the cluster level.

### Assess whether Alpha Power Increases in Response to Prolonged Pain

An identical set of analyses to those used for PAF were performed for estimates of AP.

### Assess Whether PAF and Alpha Power Changes are Scaled by Pain Intensity

PAF changes (ΔPAF) were calculated as:

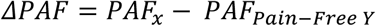

Where PAF_x_ refers to the PAF estimate at session X and Pain-Free Y refers to either Pain-Free 1 or Pain-Free 2 depending on which is most proximate to the EEG session in question (for example, Pain-Free 2 is used for CHP).

AP changes (ΔAP) were calculated as:

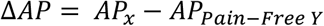

Where AP_x_ refers to the AP estimate at Session X and AP_Pain-Free Y_ refers to the AP estimate from the most temporally proximate, preceding Pain-Free session (i.e. the CHP session is most immediately preceded by the Pain-Free 2 session).

PHP pain scores were modelled at each EEG channel with a linear mixed effects model including fixed effects terms for ΔPAF, ΔAP, Visit, and the ΔPAF x ΔAP interaction as well as a random effects term including unique intercepts and slopes for ΔPAF, ΔPAF, and ΔPAF x ΔAP at each visit. CHP pain scores were modelled at each EEG channel with a linear mixed effects model including fixed effects terms for ΔPAF, ΔAP, Visit, and the ΔPAF x ΔAP interaction as well as two random effects terms (one for session and one for visit) that included unique intercepts and slopes for ΔPAF, ΔAP, and ΔPAF x ΔAP. Models were only supplied with estimates from sessions in which pain was present (PHP: PHP_on_; CHP: CHP & CHP Rekindle), and an initial cluster forming threshold of p <.0125 (4 predictors) was used. Cluster based analyses were performed separately for each predictor.

### Assess How Prolonged Pain Modulates “Fast” and “Slow” Alpha Power

Although often considered in isolation of one another, changes in PAF can be deterministically related to changes in alpha power. For example, PAF slowing can be produced via either focal power reductions in “faster” (i.e. 10-12 Hz) AP or power enhancements in “slower” (i.e. 8-12 Hz) AP. To understand how PAF changes may result from AP changes, the average 2-50 Hz (.2 Hz bins) power spectra during each EEG session was separately calculated for each EEG channel. Prior to averaging across participants at the channel level, spectra were Z-scored to account for individual differences in total alpha power amount.

For PHP, PHP_On_ spectra were subtracted from spectra obtained during Pain-Free 1. For CHP, spectra obtained during CHP and CHP Rekindle were subtracted from Pain-Free 2 spectra. To evaluate how spectral changes are related to an individual’s pain-free PAF we performed Spearman correlations to evaluate the relationship between spectral changes at each .2 Hz frequency element within the 8-12 Hz range (i.e. 8 Hz, 8.2 Hz, etc.) and PAF recorded during Pain-Free 1.

## Results

The total number of EEG datasets available at each Visit are listed in Table 1.

On average, both PHP and CHP produced moderate pain with relatively large between subject variance at both Visit 1, PHP: mean (S.D.) 26.31 (17.40); CHP: 18.81 (22.59); Rekindle: 21.86 (23.78), and Visit 2, PHP: 26.95 (20.41); CHP: mean= 20.49 (24.74); Rekindle: 20.54 (20.54). Further details regarding pain rating data can be found in Furman et al., 2020.

### PAF is Reliably Slowed at a Left, Frontocentral Cluster During Prolonged Pain

Across the entire montage, PHP Session-level effects (see *Statistics* for details) were maximally observed across a left frontocentral cluster, 8 channels, Average PHP_On_ *t* = −3.16, range = [−2.50 −3.57], summed *t* =-25.24, 97.5^th^ % null summed *t* = −5.27; Average PHP_On_ vs. PHP_Off_ & Pain-Free 2 Contrast *F* = 5.72, range: [3.92 7.72], summed *F* = 45.77, 97.5^th^ % null summed *F* = 18.66. Across this cluster, the average Cohen’s *f*^2^ was .06, range = [.04 .09]. For descriptive purposes, we present session-level estimates from the peak sensor, C3 (Figure 2C). Estimates from the posterior channel CPz that surpassed the initial cluster formation threshold can be seen in Supplementary Figure 1B. As can be seen, Peak Alpha Frequency (PAF) recorded from the left frontocentral cluster transiently slowed during PHP_On_ before returning to Pain-Free 1 levels during PHP_Off_ and Pain-free 2. In comparison, PAF at the CPz sensor was elevated during PHP_on_ before returning to baseline levels. Visit-level effects were greatest over a right posterior cluster, 8 EEG sensors, Average Visit *t* = −2.50, range = [−2.22 −2.86], summed *t* = −19.96, 97.5^th^ % null summed *t* = −5.27, and a right frontal cluster, 4 EEG sensors, Average PHP_On_ *t* = −3.16, range = [−2.50 − 3.57], summed *t* = −25.24, 97.5^th^ % null summed *t* = −17.26, which reflected that PAF was slower during Visit 2 (Figure 2B & Supplementary Figure 1A). Importantly, we could not identify any EEG channels where the expanded mixed effects model including a Session X Visit term provided a better fit to the data. Consistent with our prior studies that used a wider frequency range for PAF calculation (Furman et al., 2020), performing these analyses with 8-12 Hz PAF estimates yielded qualitatively similar, albeit weaker, results (Supplementary Figure 2) and PAF changes during PHP_On_ were not correlated with participant slider activity (Supplementary Figure 3).

**Figure 2.**
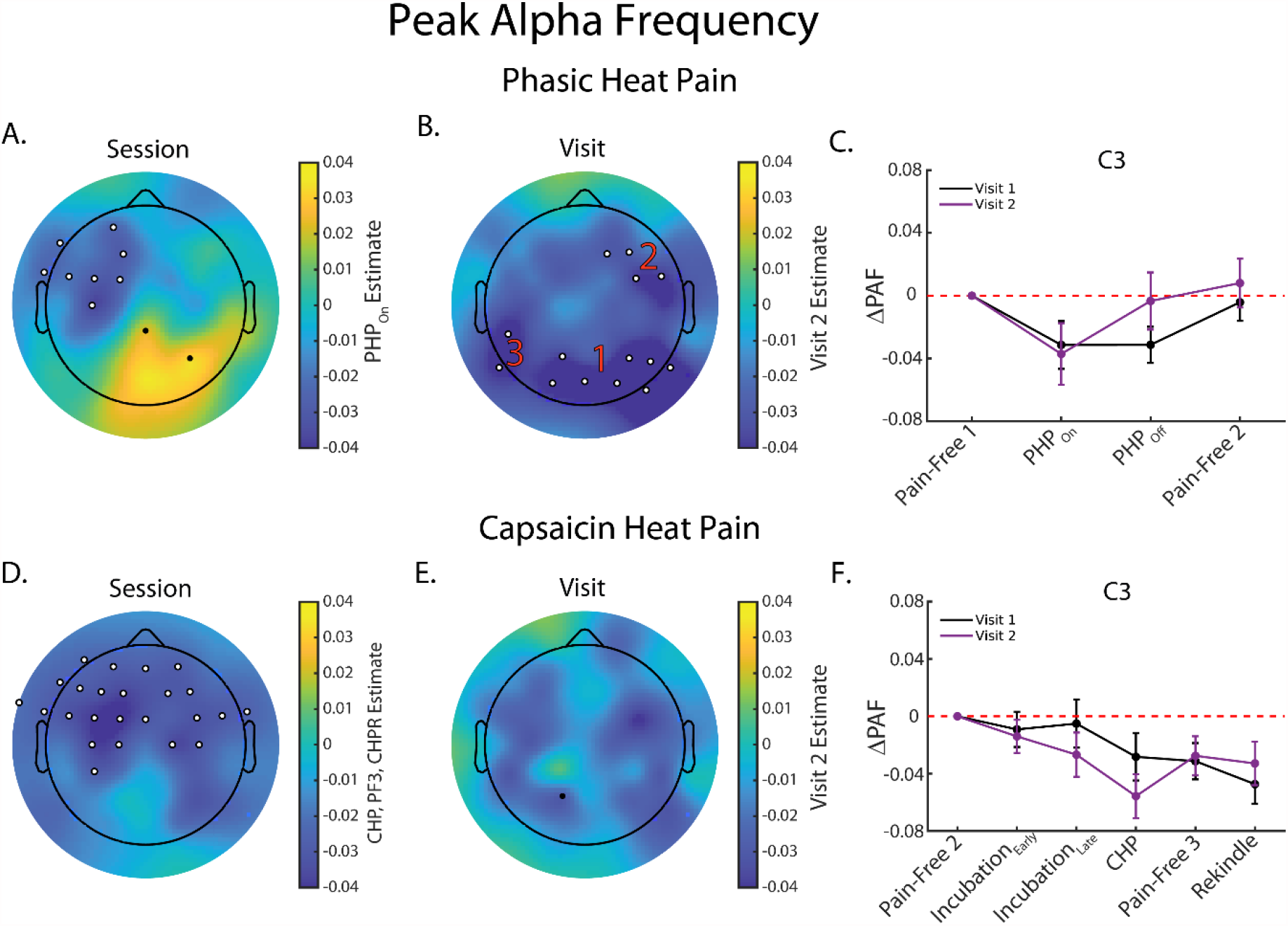
Peak Alpha Frequency (PAF) is slowed during two models of Prolonged Pain. **A**. Topoplot of the estimated effect of PHP_On_ on PAF. Cooler colors reflect that PAF decreased from Pain-free 1 to PHP_On_ while warmer colors indicate that PAF became faster. PHP_On_ PAF was found to be significantly slower than PAF recorded during Pain-Free 1 and from PAF recorded during PHP_Off_ and Pain-free 2 with this effect being observed maximally at a left, frontocentral cluster (black dots with white outlines). **B**. Topoplot of the estimated effect of Visit 2 on PAF during PHP. Cooler colors reflect that PAF was slower during Visit 2 than during Visit 1. Visit 2 PAF was found to be significantly slower, which was maximally observed over (1) and left (3) occipital clusters as well as a right frontal cluster. **C**. Line plot (mean +/- SEM) depicting PAF estimates recorded from the C3 channel during Phasic Heat Pain. For ease of visualization, PAF estimates are presented as relative to Pain-Free 1. **D**. Topoplot of the average estimated effect of CHP, Pain-Free 3, and CHP Rekindle on PAF. PAF was found to be significantly slower during all three sessions at a bilateral maximally observed over a frontocentral cluster. **E**. Topoplot of the estimated effect of Visit 2 on PAF during CHP. No significant clusters associated with the effect of Visit were identified. **F**. Line plot (mean +/- SEM) depicting PAF estimates recorded from the C3 channel during Capsaicin Heat Pain. Estimates are shown as relative to Pain-Free 2. White dots outlined with black reflect channels belonging to a significant cluster. Red numbers are used to denote instances where multiple, significant clusters were identified. If no numbers are present within a panel, all outlined channels belong to the same cluster.

For CHP, we could not identify any significant clusters for either the effect of Session or Visit. Inspecting session-level data revealed that this was likely due to sustained PAF slowing during Pain-Free 3 (Supplementary Figure 4). To identify channels demonstrating persistent PAF slowing, we performed a series of *post-hoc* contrasts on sessions occurring during the last ∼20 minutes of capsaicin exposure (CHP, Pain-Free 3, and CHP Rekindle Sessions). This revealed a large cluster of frontocentral channels (Figure 2D) where PAF slowing was greatest during the last three sessions capsaicin exposure (p < .0167 for all 3 contrasts), 24 channels, Average CHP *t* = −3.41, range = [−2.14 −4.92], summed *t* = −81.94, 97.5^th^ % null summed *t* = −10.03; Average Pain-Free 3 *t* = −3.62, range: [−2.90 −4.73], summed *t* = −86.76, 97.5^th^ % null summed *t* = −10.15; Average CHP Rekindle *t* = −3.87, range = [−2.62 −6.11], summed *t* = −92.77, 97.5^th^ % null summed *t*= −9.63. Across this cluster, the average Cohen’s *f*^2^ = .06, range = [.04 .13]. Similar magnitudes of slowing were not evident at posterior sensors and, in some instances, PAF speeding was apparent during CHP Rekindle (data now shown). The effect of Visit was not associated with any significant clusters (Figure 2E) and we could not identify any channels where the Session X Visit interaction model provided a better fit of the data. Reanalyzing the data using the expanded 8-12 Hz PAF yielded similar results (Supplementary Figure 2) and changes in PAF occurring during either CHP or CHP Rekindle were not related to participant slider movement (Supplementary Figure 3).

PAF was consistently slowed at a left, frontocentral cluster during both PHP and CHP (Figure 4A-C). Since these channels were surprisingly ipsilateral to noxious stimulation, we examined PAF effects in a mirrored channel cluster on the right side of the head. Session effects at this cluster during were qualitatively similar, albeit weaker, to those on the left side of the head (Supplementary Figure 5).

We could not identify any clusters where PAF changes were consistently correlated with reported pain intensity during PHP and CHP (Supplementary Figures 6-9).

### Alpha Power is Reliably Decreased at a Left, Frontocentral Cluster During Prolonged Pain

Significant effects of PHP Session on Alpha Power (AP; Figure 3A) were identified in a large, bilateral frontocentral cluster, 24 channels, Average PHP_On_ *t* = −4.28, range = [−2.27 −7.25], summed *t* = −102.77, 97.5^th^ % null summed *t* = −9.69; Average PHP_On_ – PHP_Off_ /Pain-Free 2 Contrast *F* = 54.30, range = [27.50 94.59], summed *t* = 1,303.10, 97.5^th^ % null summed *F* = 41.18, and in a smaller posterior cluster, 2 channels, Average PHP_On_ *t* = 2.84, range = [2.84 2.85], summed *t* = 5.70, 97.5^th^ % null summed *t* = 2.68; Average PHP_On_ – PHP_Off_ /Pain-Free 2 Contrast *F* = 5.38, range = [4.32 6.44], summed *F* = 10.76, 97.5^th^ % null summed *F* = 7.66. Across this cluster, the average Cohen’s *f*^2^ was .26, range = [.12 .49]. As can be seen in Figure 3C, AP recorded from the frontocentral cluster decreased during PHP_On_ before increasing during PHP_off_ and Pain-Free 2. In comparison, posterior AP increased throughout PHP_On_, PHP_Off_, and Pain-Free-2 (Supplementary Figure 1E). Significant effects of Visit were identified at two separate posterior clusters on the right, Average Visit *t* = −2.80, range = [−2.58 −2.95], summed *t* = −16.82, 97.5^th^ % null summed *t* = −6.07, and left sides of the head, Average Visit *t* = −2.47, range = [−2.12 −3.14], summed *t* = − 9.88, 97.5^th^ % null summed *t* = −5.82. For both clusters, the effect of Visit indicated that alpha power was reduced during Visit 2 (Figure 3B & Supplementary Figure 1D). AP estimates from several of these posterior sites, but none of the frontocentral clusters, were better captured by a linear mixed effects model including the Visit X Session interaction. This interaction always reflected that AP increases were greater at one or more Visit 2 sessions.

**Figure 3.**
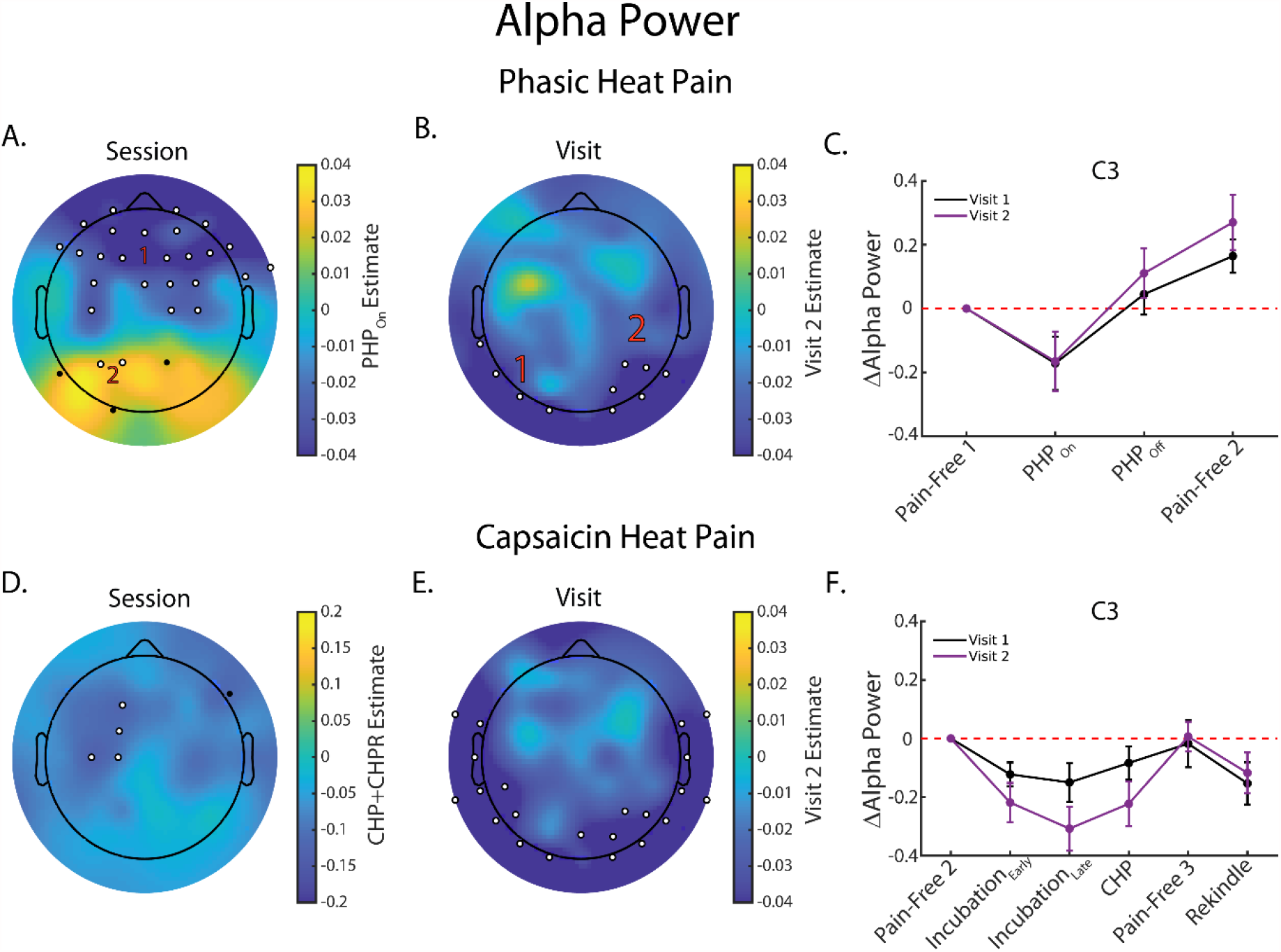
Alpha Power (AP) is decreased during two models of Prolonged Pain. **A**. Topoplot of the estimated effect of PHP_On_ on AP. PHP_On_ AP was found to be significantly less than Pain-Free 1 AP and significantly less than PHP_Off_ and Pain-free 2 AP, here maximally observed over a bilateral, frontocentral cluster (1). In contrast, AP was significantly enhanced at a left, parietal cluster (2) during PHP_On_ relative to Pain-Free 1 and relative to PHP_Off_ and Pain-Free 2. **B**. Topoplot of the estimated effect of Visit 2 on PHP AP. PHP AP was found to be significantly smaller during Visit 2 than during Visit 1 at a left (1) and right (2) posterior cluster. **C**. Line plot (mean +/- SEM) depicting AP estimates recorded from the C3 channel during Phasic Heat Pain. For ease of visualization, estimates are presented as relative to Pain-Free 1. **D**. Topoplot of the average estimated effect of CHP and CHP Rekindle on AP. AP was significantly decreased during CHP and CHP Rekindle at a left, frontocentral cluster. **E**. Topoplot of the estimated effect of Visit 2 on PHP AP. AP was significantly decreased during Visit 2 at a cluster extending over posterior and temporal channels on both sides of the head. **F**. Line plot (mean +/- SEM) depicting AP estimates recorded from the C3 channel during Capsaicin Heat Pain. Estimates are shown as relative to Pain-Free 2. For all topoplots, black dots reflect sensor-level effects surpassing the initial cluster formation threshold but not belonging to a significant cluster (see *Statistics* section for specific details on cluster formation for PHP and CHP analyses). White dots outlined with black reflect channels belonging to a significant cluster. Red numbers are used to denote instances where multiple, significant clusters were identified. If no numbers are present within a panel, all outlined channels belong to the same cluster.

AP decreased at a left, frontocentral cluster (Figure 3D) during the CHP and CHP rekindle sessions (Figure 3F), 4 channels, Average CHP *t* = −3.10, range = [−2.15 −4.44], summed *t* = −12.40; 97.5^th^ % null summed *t* = −3.92; Average Pain-Free_3_ – CHP Rekindle Contrast *F* = 5.60, range = [4.49 7.01], summed *F* = 22.38, 97.5^th^ % null summed *F* = 11.75. The average Cohen’s *f*^2^ for this cluster was .12, range = [.09 .16]. Across a large, bilateral temporo-posterior cluster (Figure 3E), AP was reduced during Visit 2, 22 channels, Average Visit *t* = −2.46, range= [−1.97 −3.30], summed *t* = −54.10, 97.5^th^ % null summed *t* = − 18.90. Data from a few channels were better described by an expanded model including the Session X Visit interaction but none belonged to the left, frontocentral cluster demonstrating significant Session effects.

Common to both PHP and CHP analyses were two left, frontocentral channels that had also been identified for the PAF conjunction analysis (Figure 4D-F). Effects at these sites were mirrored, but weaker, at right, frontocentral channels (Supplementary Figure 5). As with PAF, we could not identify any clusters demonstrating a reliable relationship between AP changes and reported pain intensity during PHP and CHP (Supplementary Figures 6-9).

**Figure 4.**
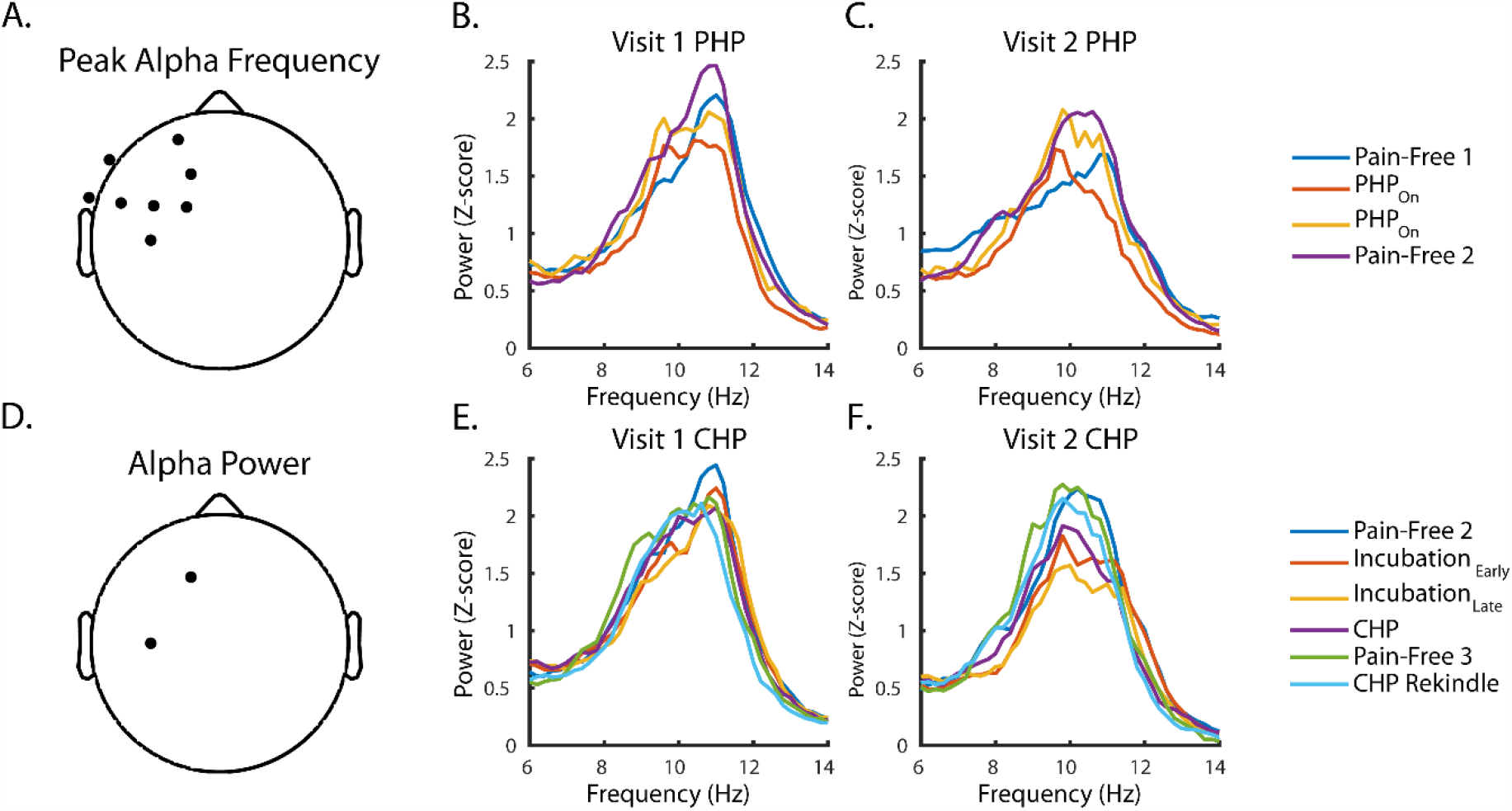
Peak Alpha Frequency (PAF) is consistently slowed and Alpha Power (AP) is reliably decreased during both Phasic Heat Pain (PHP) and Capsaicin Heat Pain (CHP). **A**. Channels denoted with a black dot in the topoplot contributed to significant clusters for both PHP and CHP PAF analyses. **B – C**. Average, z-scored spectra from the C3 sensor during PHP-related sessions at Visits 1 and 2. Increases in 8-10 Hz AP and decreases in 10-12 Hz AP can be seen at both visits (compare blue and orange spectra). **D**. Channels denoted with a black dot in the topoplot contributed to significant clusters for both PHP and CHP Alpha Power analyses. **E – F**. Same as in B-C but for CHP-relation sessions. Decreases in 8-12 Hz power are evident throughout CHP and CHP Rekindle (compare dark blue spectra to purple and and light blue spectra).

### Prolonged Pain Slows PAF by Reducing “Fast” Alpha Power

As can be seen in Figure 5, prolonged pain reliably evoked AP reductions in the “fast” 10-12 Hz range. These “fast” AP reductions were largely restricted to frontal and central sites during PHP and occurred more globally during CHP. Similarly, “fast” AP reductions were largely reversed when PHP was completed (i.e. Pain-Free 2) but not when capsaicin was still present (Pain-Free 3; Supplementary Figure 10). There did not appear to be any appreciable relationship between changes in power at each individual alpha frequency (i.e. 8 Hz, 8.2 Hz, etc.) and pain intensity (Supplementary Figure 10).

**Figure 5.**
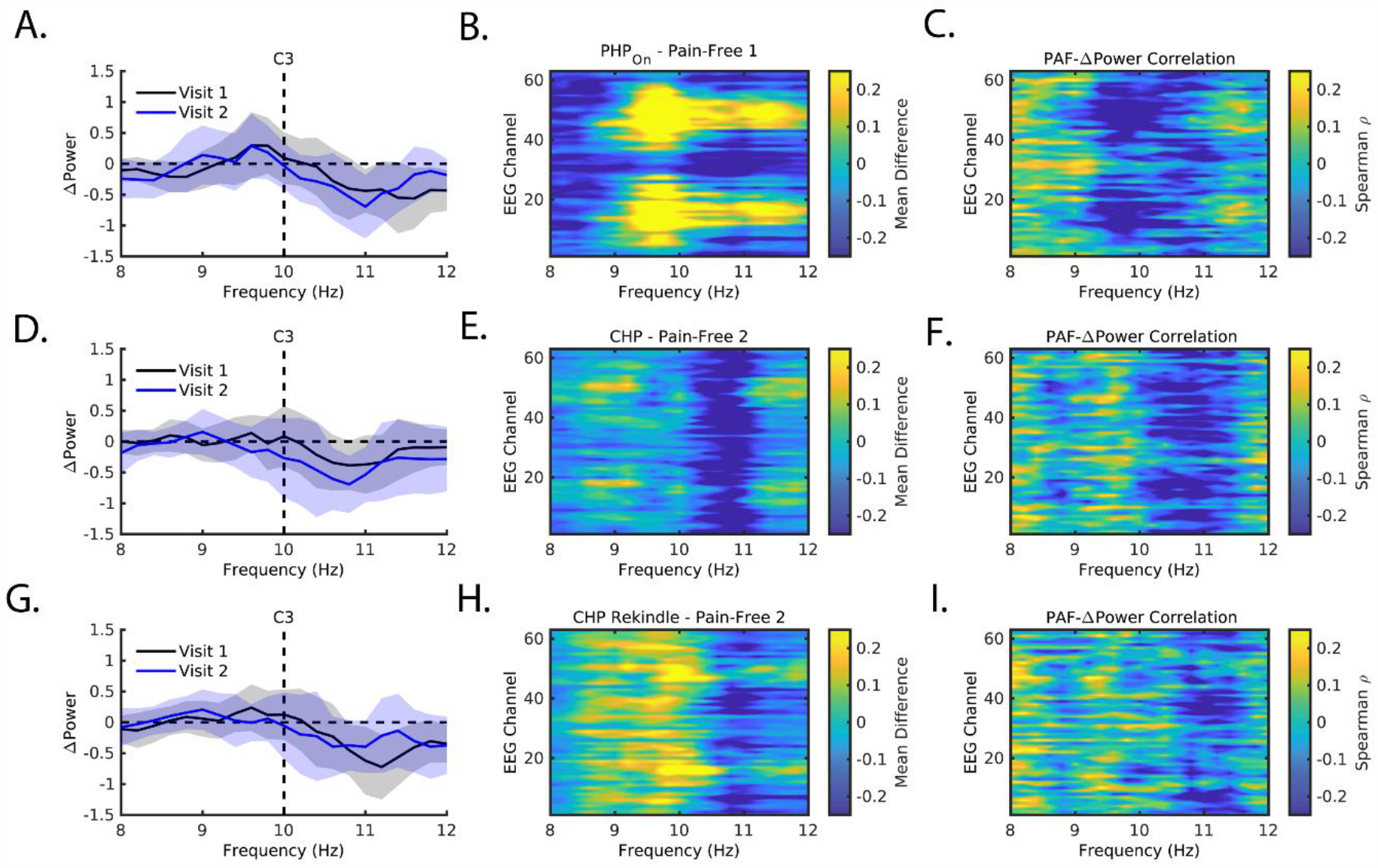
Slowing of PAF reflects focal decreases in “fast” 10-12 Hz AP during Phasic Heat Pain (A-C), Capsaicin Heat Pain (D-F), and Capsaicin Rekindle (G-I). **A**. Differences between PHP_On_ and Pain-Free 1 spectra at Visit 1 (Black) and Visit 2 (Blue). Data is taken from the C3 channel and shaded regions reflect the 95^th^ confidence interval. **B**. Same as in A., but with each row representing one of the 63 EEG channels. Increasingly warmer colors indicate greater power increases while increasingly cooler colors indicate greater power decreases. Power increases in the “fast” range are exclusive to parietal and occipital channels. **C**. Correlation between Pain-Free 1 PAF and power changes (PHP_On_ – Pain-Free 1) throughout the 8-12 Hz range for each EEG channel. Increasingly warmer colors indicate a greater positive correlation while increasingly cooler colors indicate a greater negative correlation; for ease of visualization, figures represent correlation values averaged across Visits 1 and 2. A patch of negative correlation can be seen that overlaps with the location of “fast” power decreases seen in B. **D-F**. Same as in A-C, except with spectral differences between CHP and Pain-Free 2. **G-I**. Same as in A-C, except with spectral differences between CHP Rekindle and Pain-Free 2.

Interestingly, power decreases in the “fast” range were anti-correlated with PAF. This indicated that those individual’s with relatively faster PAF experienced the greatest power decreases in this range (Figure 5 – note similarity between rows 2 and 3). To further illustrate this point, we split participants into groups based on whether their initial Pain-Free 1 PAF was above or below 10 Hz and then calculated changes to “fast” or “slow” AP for each group. As can be seen in Figure 6 for the C3 sensor, decreases in “fast” AP for individual’s with > 10 Hz PAF represented the predominant spectral change for all three prolonged pain tasks at both visits.

**Figure 6.**
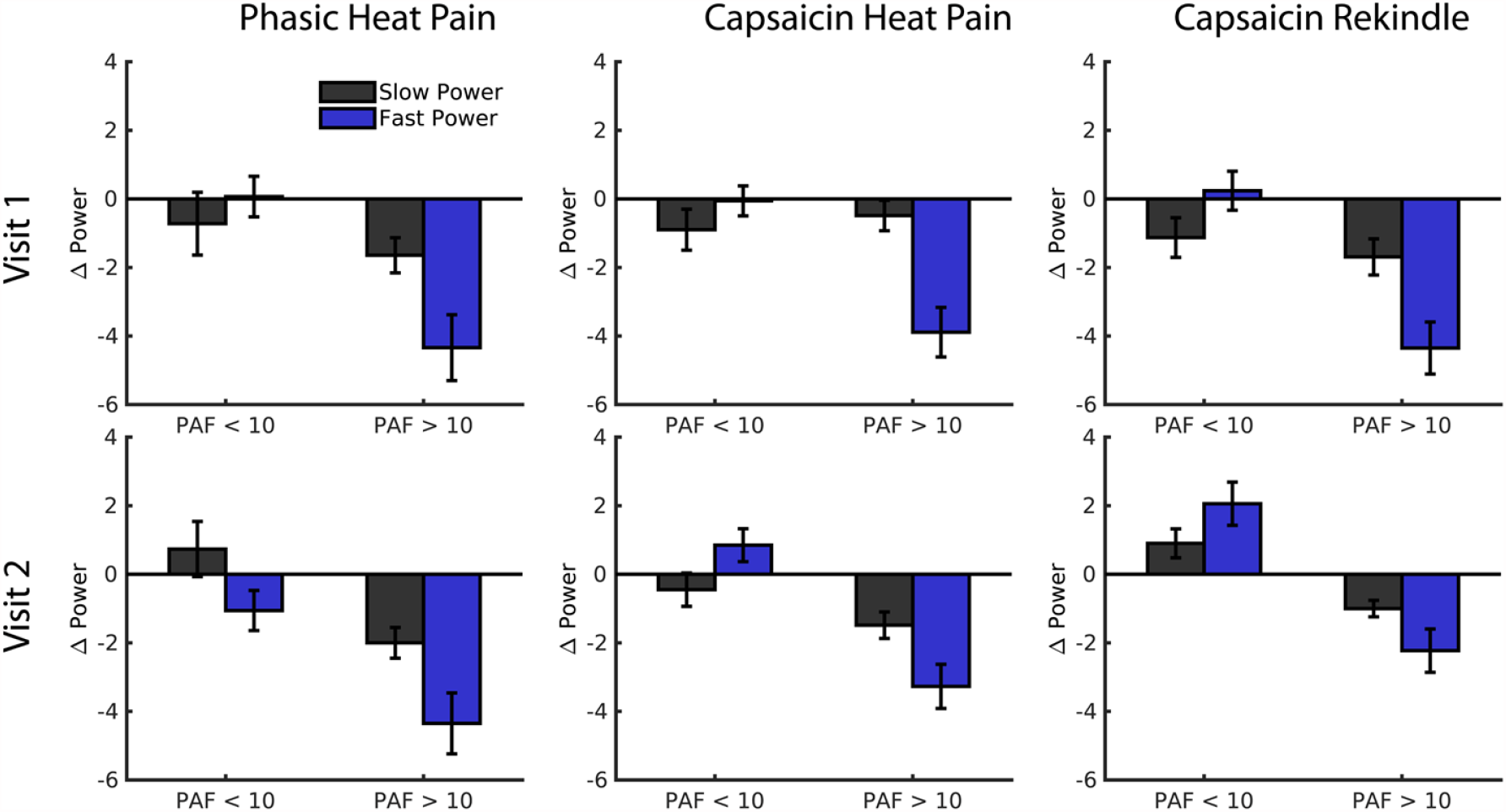
For all prolonged pain tests, decreases in “fast” 10-12 Hz AP occur predominantly in individuals with PAF greater than 10 Hz. For all panels, data from the C3 channel is shown. Participants were split based on whether their Pain-Free 1 PAF from the relevant visit was above and or below 10 Hz. Bars reflect the mean +/- 1 standard deviation.

## Discussion

The 8-12 Hz “Alpha” rhythm is hypothesized to reflect processes that control how sensory systems interact with the external environment (Wutz et al., 2018). Patients with chronic pain often present with alpha disturbances (e.g. Kim et al., 2019). Most notably, the frequency of maximal alpha power, or Peak Alpha Frequency (PAF), is slower in patients than in healthy controls. Similarly, Alpha Power (AP) decreases are associated with experimental models of acute pain while patients demonstrate elevated alpha power while at rest (Babiloni et al., 2006, Sarnthein et al., 2006). These abnormalities have been interpreted as markers of prolonged pain even though their actual coincidence with chronic pain remains unclear. Understanding if these abnormalities are evoked by prolonged pain represents an important step for potentially using these markers to identify emerging cases of chronic pain. To that end, the current study examined the reliability of PAF slowing and alpha power enhancements in response to two prolonged pain tasks, Phasic Heat Pain (PHP) and Capsaicin Heat Pain (CHP), that were repeated at two separate study visits. We report that PAF is reliably slowed and APis reliably decreased in response to prolonged pain and, furthermore, that these changes reflect a common effect specific to the “fast” (10-12 Hz) range of the alpha rhythm.

In response to PHP and CHP, PAF recorded from a cluster of left, frontocentral EEG channels was consistently slowed across Visits 1 and 2. For PHP, slowing occurred during periods where noxious temperature was applied and was restored to baseline levels both when the noxious temperature was intermittently removed during the task and during the pain-free session that followed. Perhaps more interestingly, slowing occurred throughout the late phases of CHP and even persisted when pain was reported to be absent (Pain-Free 3). As we discuss in more detail below, we believe this latter effect is unrelated to potential changes in participant alertness. Instead, we believe it represents the latent capsaicin induced sensitization that was subsequently realized during the “rekindle” phase of the protocol. If correct, this indicates that PAF slowing reflects states associated with the potential for experiencing prolonged pain (i.e. sensitization) rather than the overt experience of it. This may be particularly important for the identification of chronic pain states given the dynamic nature of its pain intensity (Jamison et al., 2001). More broadly, this is, to the best of our knowledge, the first study to demonstrate clear evidence of PAF slowing in response to prolonged pain.

Prolonged pain also reliably reduced AP at a left, frontocentral cluster that overlapped with the PAF cluster. Unlike PAF, however, AP changes were similar for both PHP and CHP with decreases occurring during periods where pain was reported (i.e. PHP_On_ and CHP Rekindle). These AP decreases subsequently returned to baseline levels when pain was absent, regardless of whether latent sensitization existed. This pattern fits well with the hypothesis that AP tracks the intensity of noxious stimuli (Nickel et al., 2017). Perhaps more importantly, prolonged pain’s failure to produce power enhancements should be taken as strong evidence that pain type (i.e. acute vs. prolonged) is not a relevant variable in shaping alpha responses in healthy individuals. The current study also demonstrates a potential mechanism by which pain-related PAF slowing is realized. More specifically, PAF changes during PHP and CHP appear to originate from focal reductions in “fast” 10-12 Hz power. In turn, these “fast” reductions leads to a relative increase in the proportion of “slow” 8-10 Hz power contributing to total AP. This process occurs to greater degrees in individuals expressing a large amount of “fast” alpha power at baseline (i.e. individuals with fast PAF). Although this might be interpreted to reflect an active pain process, our earlier work on pain-free PAF using this same dataset has identified these individuals as the least pain sensitive (Furman et al., 2020). As such, diminishment of “fast” alpha power may reflect something like a pain management resource that is depleted over time. If true, this may help to explain why individuals with slow PAF tend to be the most pain sensitive.

Importantly, reductions in “fast” alpha power were not accompanied by increases in “slow” alpha power. This result fits better with the hypothesis that the alpha rhythm is comprised of multiple oscillators rather than a single oscillator whose center frequency changes (Figure 7, Klimesch et al., 1997; Lobier et al., 2018; Benwell et al., 2019, Barry et al., 2020). In the context of pain, there appears to be at least two relevant oscillators (Egsgaard et al., 2009). Whereas the “slow” 8-10 Hz range has been associated with pain sensitivity (Nir et al., 2012) the “fast” 10-12 Hz range has been associated with pain tolerance (Furman et al., 2020). Given this apparent dichotomy, deciphering the cognitive processes represented by each oscillator as well as determining whether the two share a similar or disparate anatomical localization represents important targets for future research.

**Figure 7.**
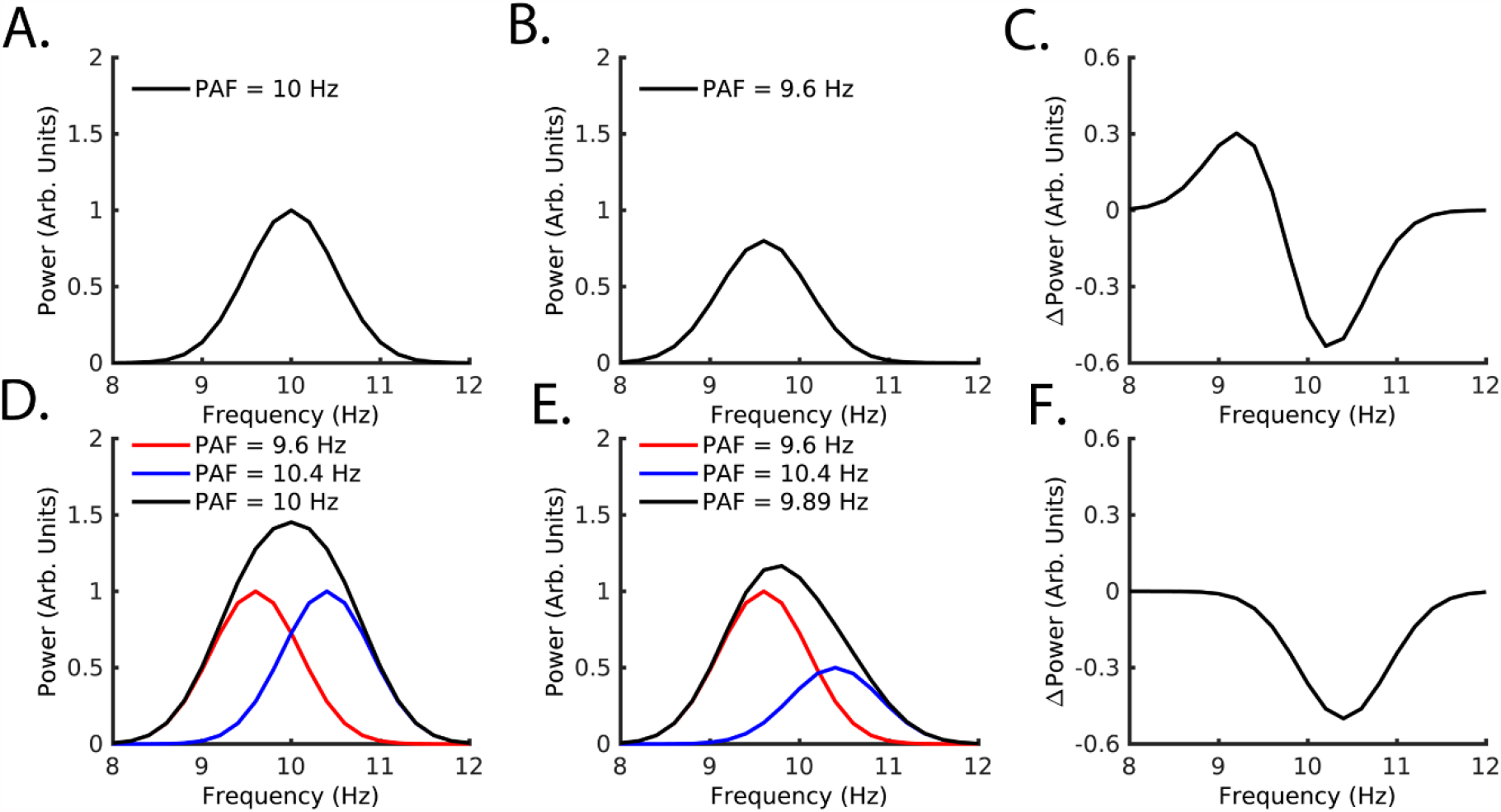
Single (**A. – C**.) and multiple (**D. – F**.) oscillator mechanisms for slowing peak alpha frequency and reducing alpha power. **A. – C**. In the case where a single oscillator (A) both reduces its speed and power (B), the pattern of differences will represent a sigmoid and the sum of differences will equal 0. **D. – F**. Two oscillators (D, red and blue denote each oscillator while the black represents their summed representation) can produce PAF and power changes without changing their frequency location (E). In this instance, spectral changes will present a focal reduction and the sum differences will not be equal to 0. Data from Figures 5 and 6 appear to more closely follow a multi-oscillator scheme.

Before discussing the potential clinical implications of these findings, we acknowledge some possible limitations of the current work. First, PAF slowing during cognitive tasks has been reported to occur passively with the passage of time and has been suggested to reflect changes in alertness and/or stimulus salience (Benwell et al., 2019). Given that our experiments occurred over the course of hours, it is possible that any identified PAF slowing may owe to these sources rather than prolonged pain. Time-associated slowing of PAF is accompanied, however, by enhancements rather than decreases in AP. This suggests that the effects of time on PAF are realized through increases in “slow” 8-10 Hz power. This appears to be an entirely different mechanism than the one identified in the current work. Second, effects of prolonged pain on PAF and AP were primarily identified at EEG channels ipsilateral to noxious stimulation and, perhaps more importantly, contralateral to the hand providing continuous ratings. We believe these effects are unlikely to reflect motor activity for two reasons: 1) Effects were present at contralateral channels, albeit at weaker strengths, and were strongly correlated with those occurring at ipsilateral sites, and 2) we could not find any evidence of a consistent relationship between putative motor activity (i.e. rating device use) and changes in either PAF or AP (Supplementary Figure 3).

These limitations aside, the current findings serve to further expand our understanding of how PAF abnormalities may relate to chronic pain. While our prior work with the very same dataset has suggested that pain-free PAF reflects traits associated with chronic pain vulnerability (i.e. pain sensitivity; Furman et al., 2020), the current work indicates that PAF slowing is a product of states associated with prolonged pain. This opens the door to the possibility that there exist two populations of people at risk for developing chronic pain: 1) Those with slow PAF before injury, and 2) those whose PAF slows after the onset of injury.

Although it remains to be seen whether chronic pain produces PAF slowing, monitoring of PAF slowing after injury could provide important real-time information to caregivers, even when overt pain is absent, about the development of states that may be associated with chronic pain. Secondly, new interventions that aim to either enhance “fast” alpha power prior to injury, especially in those individuals with slow PAF, may be useful in preventing chronic pain. Similarly, restoring “fast” AP after injury may hold promise for preventing the transition to chronic pain. Focal reductions in the “fast” alpha range may explain why previous attempts to modulate pain via non-invasive alpha modulation have not been as successful as anticipated (Arendsen et al., 2018; Ahn et al., 2019). These studies have generally modulated the alpha rhythm at a frequency, roughly 10 Hz, that is shown to respond neutrally to prolonged pain in the current study (Figure 5).

In summary, we present evidence that PAF is reliably slowed and that AP is reliably reduced during pain. Based on its occurrence during the non-painful portions of capsaicin exposure, PAF slowing appears to reflect the emergence of states that promote pain such as sensitization. In contrast, changes in AP appear to track the presence of a noxious stimulus. Importantly, we demonstrate that both effects derive from a common source: focal reductions in “fast” alpha power. Taken together, these results highlight PAF slowing as a marker of states associated with prolonged pain and suggest that manipulations of the 10-12 Hz band of the alpha rhythm may hold promise for combatting emerging cases of chronic pain.

## Acknowledgements

Patent pending (D.A.S., A.M., A.J.F.).

