## Supplemental Data for "Prolonged Pain Reliably Slows Peak Alpha Frequency by Reducing Fast Alpha Power"

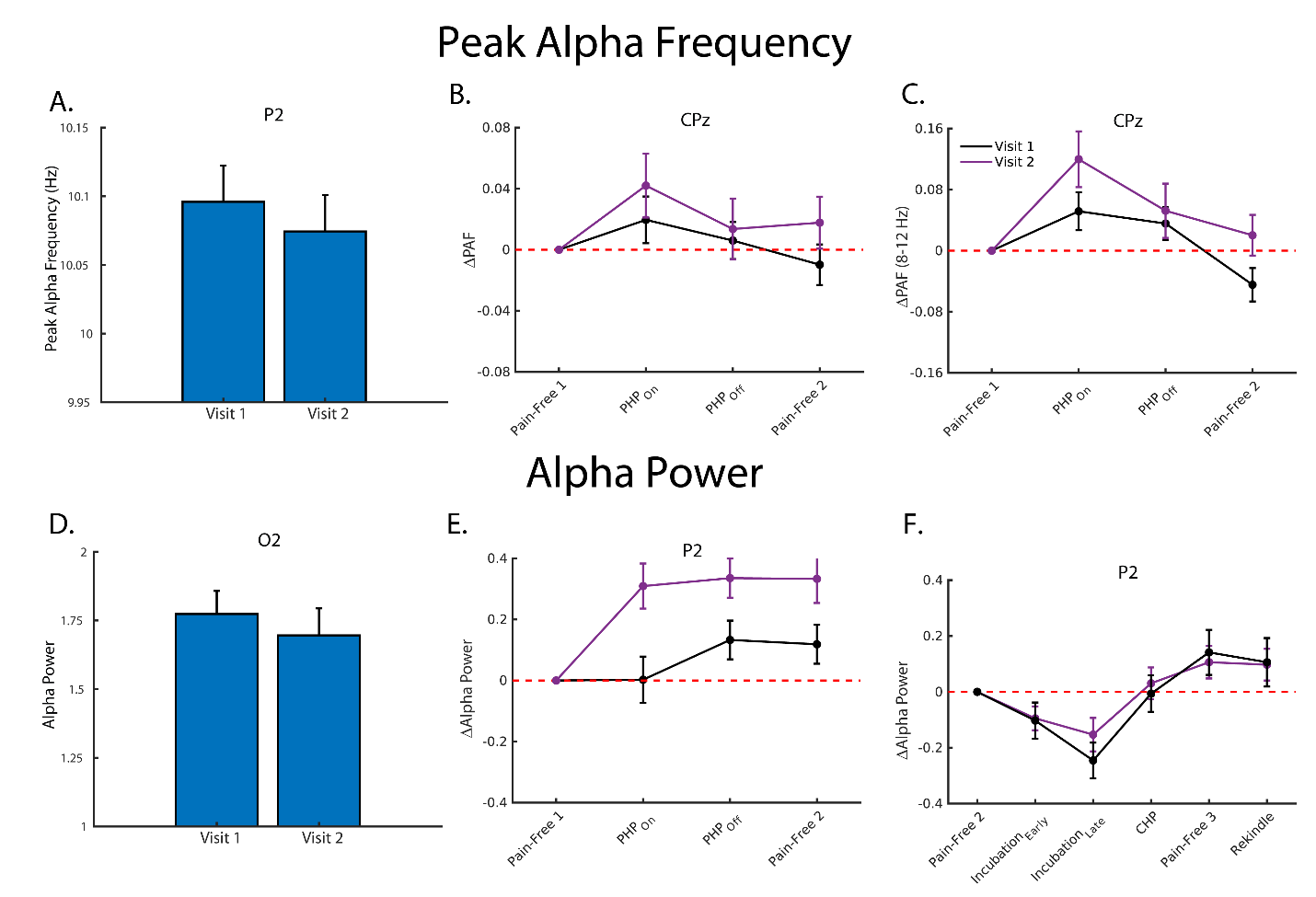

**Supplementary Figure 1. A.** Bar plots (mean +/- SEM) depict grand average Peak Alpha Frequency (PAF) estimates recorded during PHP at Visits 1 and 2. As can be seen, PAF is slower at Visit 2. Data is taken from the P2 channel, which corresponded to the largest Visit effect within the identified cluster. **B.** Line plot (mean +/- SEM) depicting PAF estimates recorded from the CPz channel during PHP. Estimates are shown as relative to Pain-Free 1. At CPz, PAF is transiently increased during PHP_On_ before returning to baseline levels during PHP_Off_ and Pain-Free 2. **C.** Same as in B., but for PAF calculated using the wider 8-12 Hz range. **D.** Bar plots (mean +/- SEM) depict grand average Alpha Power (AP) estimates during CHP at Visits 1 and 2. As can be seen, AP was reduced during Visit 2. Data is taken from the O2 channel, which corresponded to the largest Visit effect within the identified cluster. **E.** Line plot (mean +/- SEM) depicting AP estimates recorded from the P2 channel during PHP. Estimates are shown as relative to Pain-Free 1. P2 AP is increased during PHP_On_, PHP_Off,_ and during Pain-Free 2. **F.** Line plot (mean +/- SEM) depicting PAF estimates recorded from the P2 channel during CHP. Estimates are shown as relative to Pain-Free 2. At P2, AP is decreased during capsaicin incubation before returning to baseline levels during CHP and then surpassing baseline levels during Pain-Free 3 and CHP Rekindle.

*Session Effects Are Weaker but Present When Using 8-12 Hz PAF*

Data for the effect of Session and Visit on PHP PAF calculated with the extended 8-12 Hz range are presented in Supplementary Figure 2A-C. Although we were unable to identify a significant effect of Session at the previously identified left frontocentral cluster, the direction and magnitude of effects were similar, 8 channels, Average PHP_On_ *t* = -1.39, range = [-2.98 .39], summed *t* = -11.11; Average PHP_On_ vs. PHP_Off_ & Pain-Free 2 Contrast *F* = 1.09, range: [<.01 3.08], summed *F* = 8.77. These weaker results were not unexpected given that using the 8-12 Hz range likely introduces data from with non-alpha rhythms (Furman et al., 2018). Perhaps more importantly, summing the relevant *t* or *F* statistic across the cluster revealed that the cluster wide effects surpassed the 97.5^th^ percentile of both the PHP_On­_, *t* = -5.31, and PHP_On_ vs. PHP_Off_ & Pain-Free 2 Contrast, *F* = 5.49, null distributions. Additionally, we identified a large posterior cluster sensitive to the effects of Session, 24 channels, Average PHP_On_ *t* = 3.77, range = [2.14 5.32], summed *t* = 90.40, 97.5^th^ % null summed *t* = -9.48; Average PHP_On_ vs. PHP_Off_ & Pain-Free 2 Contrast *F* = 7.67, range: [4.10 16.48], summed *F* = 184.18, 97.5^th^ % null summed *F* = 39.36, which reflected that PAF recorded from these channels was increased during periods when the temperature was on (Supplementary Figure 2C). Across this cluster, the average Cohen’s *f*^2^ for the effect of Session = .10, range = [.06 .18]. As before, we identified a large posterior cluster demonstrating that PAF was slower during Visit 2, 14 channels, Average Visit *t* = -2.50, range = [-2.07 -3.12], summed *t* = -35.07, 97.5^th^ % null summed *t* = -27.16, as well as a smaller central cluster, 5 channels, Average Visit *t* = -2.31, range = [-2.00 -2.60], summed *t* = -11.53, 97.5^th^ % null summed *t* = -10.52. Finally, we could not find any instances where data from these channels were better described by a model including a term for the Session X Visit interaction.

Data for the effect of Session and Visit on CHP PAF calculated with the extended 8-12 Hz range are presented in Supplementary Figure 2D-F. For the effect of session, several channels from the originally identified cluster surpassed the initial cluster forming threshold, 7 channels, Average CHP *t* = -2.88, range = [-2.59 -3.50], summed *t* = -20.17, 97.5^th^ % null summed *t* = -5.13; Average Pain-Free 3 *t* = -4.14, range = [-3.33 -5.30], summed *t* = -29.04, 97.5^th^ % null summed *t* = -5.07, and CHP Rekindle, Average CHP Rekindle *t* = -4.60, range = [-3.87 -5.27], summed *t* = -32.19, 97.5^th^ % null summed *t* = -4.89. Across this cluster, the average Cohen’s *f*^2^ for the effect of Session = .11, range = [.09 .14]. For the entirety of the originally identified, PAF was found to be significantly slowed during all three late CHP sessions, 24 channels, Average CHP *t* = -1.78, range = [-.39 -3.50], summed *t* = -42.94, 97.5^th^ % null summed *t* = -10.49; Average Pain-Free 3 *t* = -4.20, range = [-2.16 -5.36], summed *t* = -100.78, 97.5^th^ % null summed *t* = -9.88, and CHP Rekindle, Average CHP Rekindle *t* = -3.61, range = [-1.58 -5.27], summed *t* = -86.55, 97.5^th^ % null summed *t* = -9.42. The effect of Visit was not associated with any significant clusters and no channels were better described by a model including a term for the Session X Visit interaction.

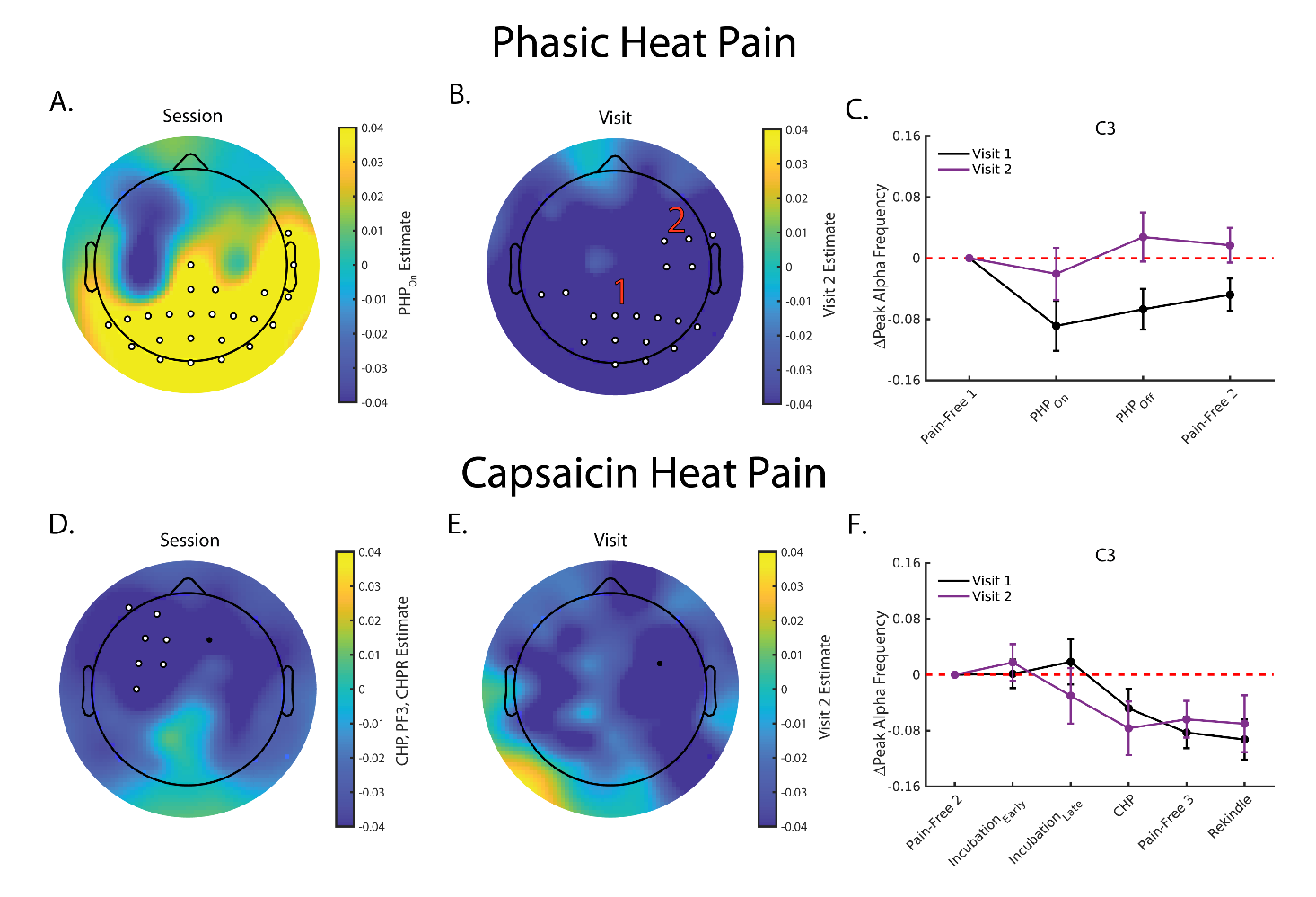

**Supplementary Figure 2**. Prolonged pain still produces PAF slowing at frontocentral channels even when PAF is calculated using the wider 8-12 Hz range. **A.** Topoplot of the estimated effect of PHP_On_ on 8-12 Hz PAF. Cooler colors reflect that PAF decreased from Pain-free 1 to PHP_On_ while warmer colors indicate that PAF became faster. PHP PAF was found to be significantly increased at a large posterior cluster. Although no significant frontocentral clusters were identified, PHP_On_ PAF was slowed at series of left frontocentral channels that overlapped with those identified in the main text. **B.** Topoplot of the estimated effect of Visit 2 on 8-12 Hz PAF during PHP. Cooler colors reflect that PAF was slower during Visit 2 than during Visit 1. Significant decreases in Visit 2 PAF were noted at a posterior (1) and central (2) cluster. **C.** Line plot (mean +/- SEM) depicting PAF estimates recorded from the C3 cannel during Phasic Heat Pain. For ease of visualization, PAF estimates are presented as relative to Pain-Free 1. As can be seen, PAF is transiently slowed to varying degrees at the two visits before beginning to return to baseline levels at PHP_Off_ and Pain-Free 2. **D.** Topoplot of the average estimated effect of CHP, Pain-Free 3, and CHP Rekindle on 8-12 Hz PAF. PAF was found to be significantly slower during all three sessions at a number of left, frontocentral belonging to the originally identified cluster. **E.** Topoplot of the estimated effect of Visit 2 on 8-12 Hz PAF during CHP. As in the original analysis, no channels in the left, frontocentral cluster demonstrated a significant effect of Visit. **F.** Line plot (mean +/- SEM) depicting 8-12 Hz PAF estimates recorded from the C3 channel during Capsaicin Heat Pain. Estimates are shown as relative to Pain-Free 2. As before, PAF became increasingly slowed throughout Capsaicin application. For all topoplots, black dots reflect sensor-level effects surpassing the initial cluster formation threshold but not belonging to a significant cluster (see Statistics section for specific details on cluster formation for PHP and CHP analyses). White dots outlined with black reflect channels belonging to a significant cluster. Red numbers are used to denote instances where multiple, significant clusters were identified. If no numbers are present within a panel, all outlined channels belong to the same cluster.

*Session Effects on PHP and CHP Cannot be Explained by Slider Movement*

Most Session effects for PHP and CHP appeared at the side of the head contralateral to the hand used to provide pain ratings. To account for the possibility that these effects reflect preparatory or motor processes rather than those related to the experience of pain, we investigated the relationship between movement of the manual analog device and changes to PAF and Alpha Power (AP). To get an idea of how much an individual moved the rating device, movement was quantified as the sum of the absolute first derivative of the z-scored pain ratings (see Supplementary Figure 3). We elected to normalize pain ratings in order to ensure that our estimate of movement was not confounded by pain. Results were nearly identical when we calculated rating device movement without normalization (data not shown). If rating device movement is responsible for the Session effects, then changes in PAF or AP should be negatively related to our measure of movement.

As can be seen in Supplementary Figure 3, no relationship reliably emerged at the frontocentral channels identified in our main analyses. Indeed, the only instance where we did detect significant correlations between slider movement and CHP Rekindle ΔPAF at these sites was in *opposite* direction to what was expected. Given these results, we feel confident that our Session level effects cannot be solely explained by processes related to slider movement.

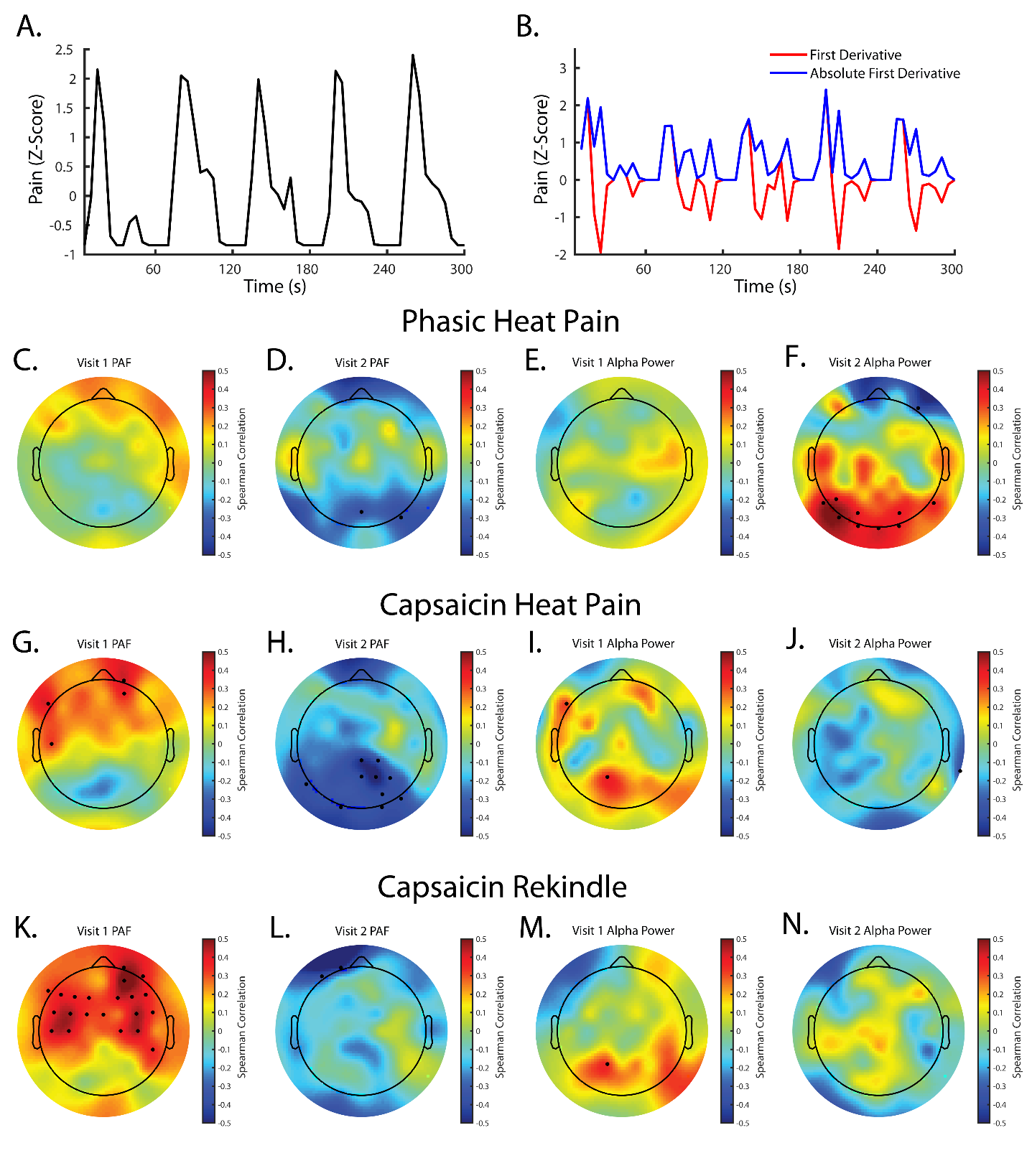

**Supplementary Figure 3.** Movement of the rating slider is not reliably related to PAF or Alpha Power (AP) changes. **A.** An example of z-transformed pain rating data from the PHP task. **B.** The first derivative and absolute first derivative of the pain rating data in A. The summed absolute first derivative was used as a proxy for slider movement since it reflects any change in position of the slider. **C-D**. Topoplots reflect the correlation between our metric for slider movement and changes in PHP PAF or AP (PHP_On_ – Pain-Free 1) at Visit 1 or Visit 2. Black dots reflect channels demonstrating a correlation with an associated p value < .05. No apparent relationship exists between slider movement and PAF or AP changes at frontocentral channels. **G-J.** Topoplots reflect the correlation between our metric for slider movement and changes in PHP PAF or AP (CHP – Pain-Free 2) at Visit 1 or Visit 2. None of the identified frontocentral channels from the main text show a relationship between spectral changes and slider movement. **K-L**. Topoplots reflect the correlation between our metric for slider movement and changes in CHP Rekindle PAF or AP (CHP Rekindle – Pain-Free 2) at Visit 1 or Visit 2. Only during Visit 1 is there a relationship between changes in PAF and slider movement. This relationship is in the opposite direction to what would be expected as increased slider movement is associated with greater PAF speeding. For all topoplots, black dots reflect channels demonstrating a relationship between PAF and slider movement with an associated p < .05.

*Peak Alpha Frequency is slowed during Pain-Free 3 and Alpha Power is elevated during Pain-Free 2 and 3*

Across a mostly overlapping, widespread cluster of channels, Pain-Free 3 PAF was slowed in comparison to either to Pain-Free 1, 54 channels, average *t* = -4.00, range = [-2.47 -6.07], summed *t* = -215.83, 97.5^th^ % null summed *t* = -14.32, and Pain-Free 2, 45 channels, average *F* = 11.90, range = [6.01 26.21], summed *F* = 535.72, 97.5^th^ % null summed *F* = 65.61, Figure 3A. An example of this latter effect taken from the C3 channel can be seen in Figure 4B. We were unable to identify any clusters where PAF was significantly different between Pain-Free 2 and Pain-Free 1 and, furthermore, could only identify a single channel, CP5, that surpassed our initial cluster forming threshold. We could not find any evidence that the Session X Visit interaction expanded model explained more variance at any one of these sensors, indicating that these effects were stable across Visits. We obtained nearly identical results when using PAF estimates calculated with the wider 8-12 Hz range (Supplementary Figure 3A). As we discuss below, the apparent slowing of PAF during Pain-Free 3 appears to be a result of the CHP model specifically, given that similar effects were not noted during PHP, and led us to replace the *a priori* CHP contrasts with more suitable post-hoc comparisons.

In comparison to Pain-Free 1, AP was elevated across almost the entire montage during both Pain-Free 2, 35 channels, average *t* = 3.42, range = [2.52 6.12], summed *t* = 64.80, 97.5^th^ % null summed *t* = 11.32, and Pain-Free 3, average *t* = 4.19, range = [2.58 7.81], summed *t* = 150.49, 97.5^th^ % null summed *t* = 11.38. An example of this effect taken from the C3 channel is shown in Figure 5. We further identified that AP was greater during Pain-Free 3 than during Pain-Free 2 at a pair of clusters located over right frontal, 2 channels, average F = 6.92, range = [6.71 7.128], summed *F* = 13.84, 97.5^th^ % null summed *F* = 7.66, and right posterior sensors, 4 channels, average *F* = 9.21, range = [6.70 12.38], summed *F* = 36.84, 97.5^th^ % null summed *F* = 10.94, Figure 2B. The effect of Visit was significant at a pair of posterior clusters, left cluster: 4 channels, average *t* = -2.32, range = [-2.10 -2.90], summed *t* = -9.29, 97.5^th^ % null summed *t* = -5.82; right cluster: 6 channels, average *t* = -2.69, range = [-2.28 -2.92], summed *t* = -16.14, 97.5^th^ % null summed *t* = -5.54, indicating that AP was relatively lower at Visit 2. Importantly, we could only identify a single sensor, O1, where the Session X Visit expanded model explained more variance in AP. Thus, it these AP increases are reliable effects across Visits. Despite these differences between Pain-Free sessions, it is clear from the data that these differences represent baseline “resets”. That is, subsequent AP changes are in relation to the most recent Pain-Free session rather than Pain-Free 1 (Supplementary Figure 3B). As such, no revision of our *a priori* AP analyses are necessary.

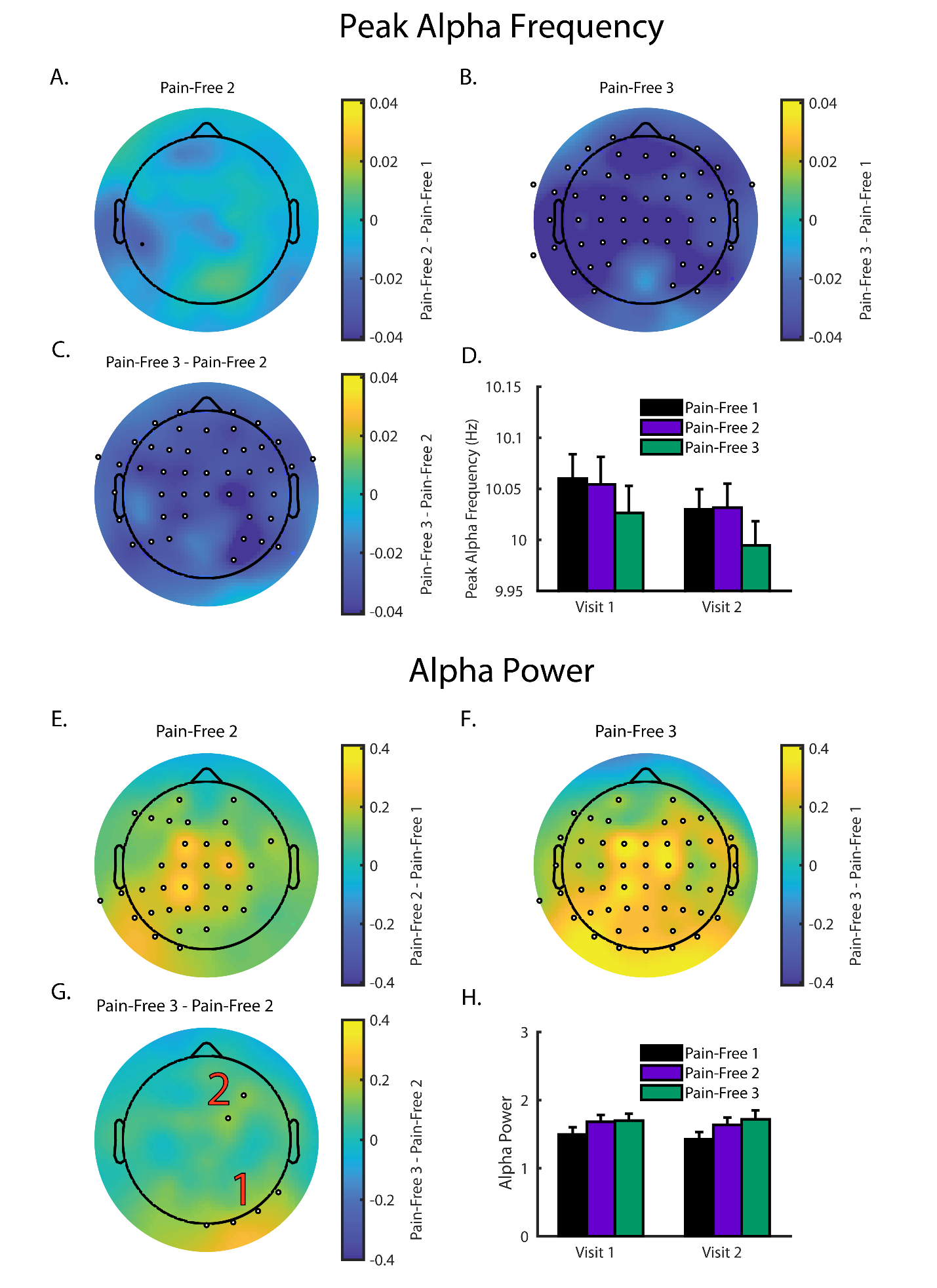

**Supplementary Figure 4.** Peak Alpha Frequency (PAF) is decreased during Pain-Free 3 while Alpha Power (AP) is elevated during Pain-Free 2 and Pain-Free 3. **A.** Topoplot of the estimated effect of Pain-Free 2 on PAF. Cooler colors reflect that PAF decreased from Pain-Free 1 to Pain-Free 2 while warmer colors indicate that PAF became faster. PAF appeared to remain relatively stable between the two pain-free sessions. **B.** Topoplot of the estimated effect of Pain-Free 3 on PAF. Pain-Free 3 PAF was significantly slower at a large number of channels. **C.** Topoplot demonstrating the comparison between Pain-Free 3 and Pain-Free 2 PAF. PAF was significantly slower at a large number of channels during Pain-Free 3. **D.** Bar plots (mean +/- SEM) demonstrating pain-free PAF estimates from the C3 channel. PAF is reduced during Pain-Free 3 at both visits. **E – G.** Same as in A-D but using AP. In comparison to Pain-free 1, AP is significantly elevated at a wide range of channels during Pain-Free 2 and Pain-Free 3. AP is relatively similar, however, between Pain-Free 2 and Pain-Free 3. **H.** Bar plots (mean +/- SEM) demonstrating pain-free AP estimates from the C3 channel. AP is enhanced during Pain-Free 2 and Pain-Free 3 at both visits. For all topoplots, black dots reflect sensor-level effects surpassing the initial cluster formation threshold but not belonging to a significant cluster (see Statistics section for specific details on cluster formation for PHP and CHP analyses). White dots outlined with black reflect channels belonging to a significant cluster. Red numbers are used to denote instances where multiple, significant clusters were identified. If no numbers are present within a panel, all outlined channels belong to the same cluster.

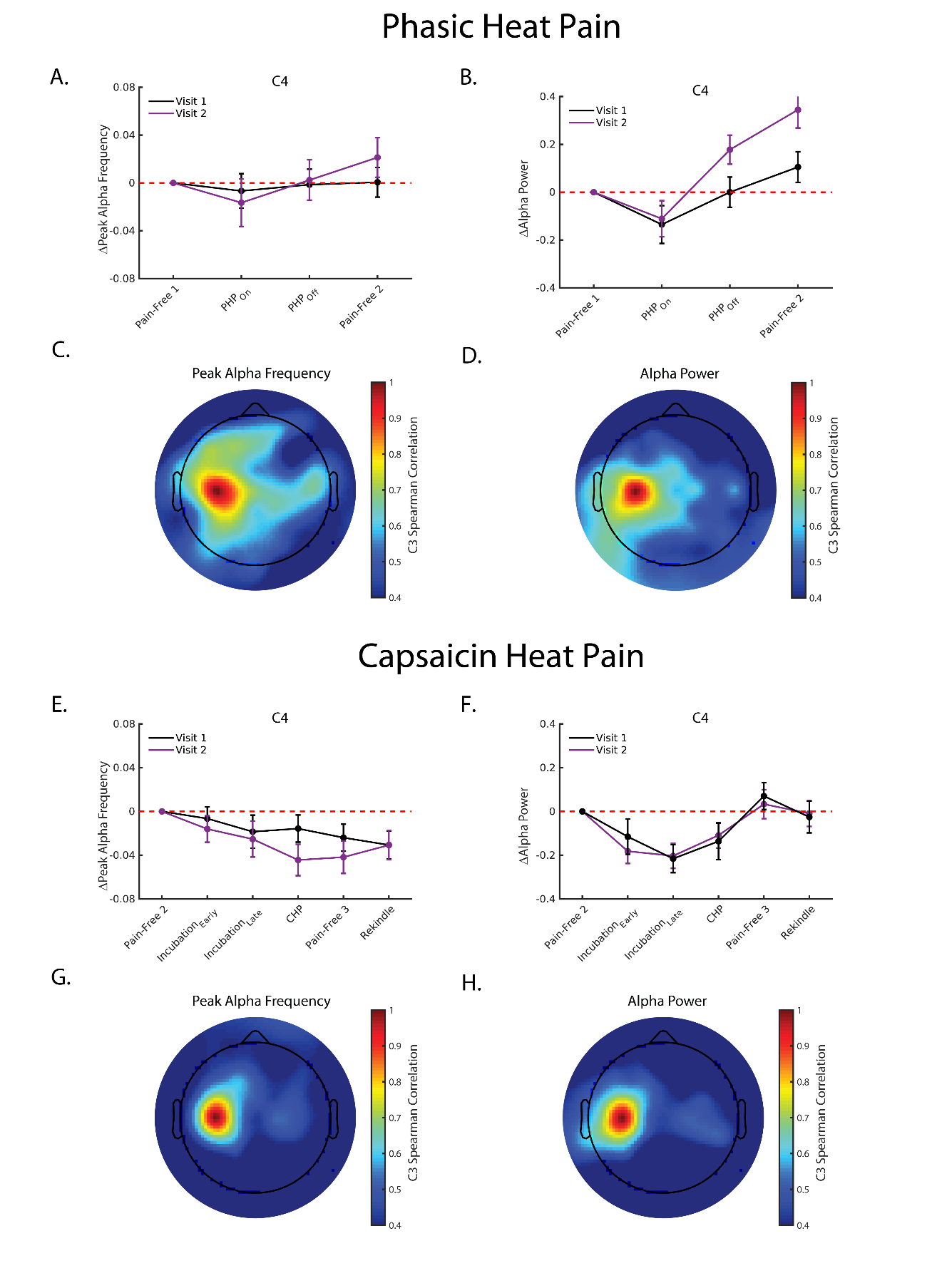

**Supplementary Figure 5.** Effects of Prolonged Pain on PAF and Alpha Power (AP) at right, frontocentral channels are similar to those seen at left frontocentral channels. **A.** Line plot (mean +/- SEM) depicting PAF estimates recorded from the C4 channel during Phasic Heat Pain. For ease of visualization, PAF estimates are presented as relative to Pain-Free 1. In comparison to the C3 channel (Figure 2C), C4 PAF undergoes less slowing during PHP_on_ before returning to baseline levels during PHP_Off_ and Pain-Free 2. **B.** Line plot (mean +/- SEM) depicting AP estimates recorded from the C4 channel during Phasic Heat Pain; estimates are presented as relative to Pain-Free 1. As at the C3 channel, AP is transiently reduced during PHP_On_. **C.** Topoplot reflects the correlation between estimates of C3 PHP ΔPAF (PHP_On_ – Pain-Free 1) and estimates of PHP ΔPAF at every other channel. Warmer colors reflect greater similarity between estimates from that channel and the C3 channel. Strong similarity is seen between right, central channels (i.e. C2, C4) and the C3 channel. **D.** Same as in C but using PHP ΔAP. Similarity between the C3 sensors and right, central channels is once again seen. **E.** Line plot (mean +/- SEM) depicting PAF estimates, shown relative to Pain-Free 2, recorded from the C4 channel during Phasic Heat Pain. Similar to what is seen at the C3 channel, C4 PAF slows throughout capsaicin exposure. **F.** Line plot (mean +/- SEM) depicting AP estimates recorded from the C4 channel during Phasic Heat Pain; estimates are presented as relative to Pain-Free 2. AP decreases throughout the initial capsaicin exposure period (Incubation_Early_ **–** CHP) but does not decrease during CHP Rekindle. **G.** Same as in C but for CHP ΔPAF (CHP – Pain-Free 2). Similarity is once again seen between estimates from the C3 channel and mirrored channels on the right side of the head. **H.** Same as in G but using CHP AP estimates. C3 estimates are similar to those seen at right, central channels.

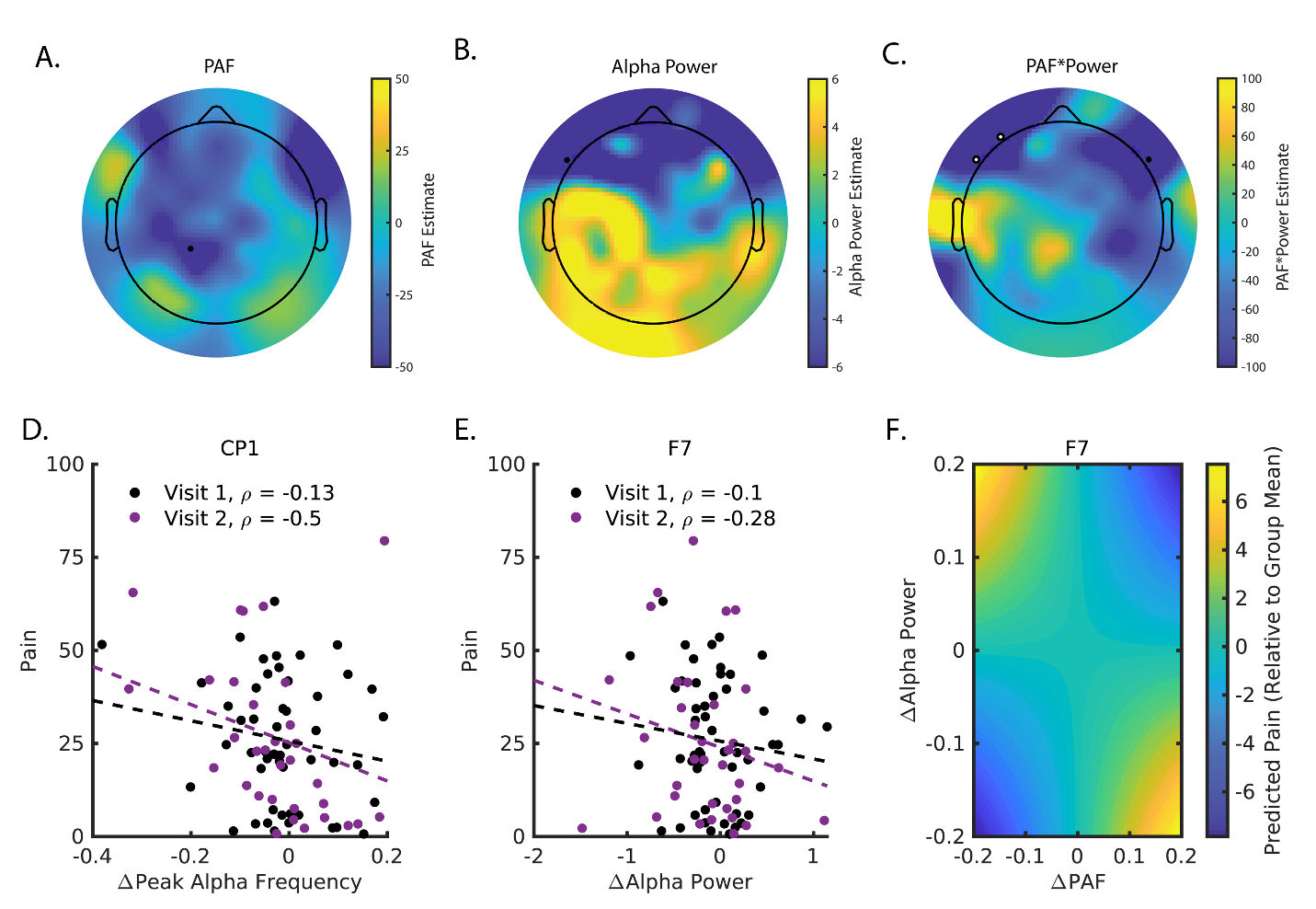

**Supplementary Figure 6.** Changes in PAF (ΔPAF) and Alpha Power (ΔAlpha Power) during PHP are not scaled by pain intensity. No significant clusters were associated with either changes in PAF or Alpha Power (**A. & B.**). Scatter plots in **D. & E.** reflect simple correlations between PHP_On_ pain intensity and spectral changes taken from channels surpassing initial cluster forming threshold. **C.** Pain intensity was related to the ΔPAF*ΔAlpha Power interaction at a right, frontocentral cluster. As shown in **F.**, slowing of PAF with concurrent increases in Alpha Power was associated with greater pain while PAF speeding and power increases were associated with less pain. Data was generated by multiplying hypothetical changes to ΔPAF and ΔAlpha Power with the interaction β obtained from the linear mixed model described in the text. For all topoplots, black dots reflect sensor-level effects surpassing the initial cluster formation threshold (p < .017). Black dots with white centers reflect channels belonging to a significant cluster.

*Prolonged Pain Intensity is not Reliably Scaled by Changes in PAF or Alpha Power*

For PHP (Supplementary Figure 6), a linear mixed model identified that the interaction between changes in PAF and changes in Alpha Power (ΔPAF x ΔAP) at a left, frontal cluster were predictive of pain intensity, 2 channels, average ΔPAF x ΔAP *t* = -3.08, range = [-3.06 -3.10], summed *t* = -6.17, 97.5^th^ % null summed *t* = -2.84, which reflected that greater pain was associated with either AP decreases and PAF speeding or AP increases and PAF slowing. On average, the Cohen’s *f*^2^ for ΔPAF x ΔAP at these sensors was .10, range = [.10 .11]. No significant clusters were associated with either changes in ΔPAF or ΔAP.

The intensity of CHP and CHP Rekindle pain were negatively associated with ΔAP at a left frontal cluster (Supplementary Figure 7, 3 channels, average ΔAP *t* = -2.89, range = [-2.43 -3.31], summed *t* = -8.66, 97.5^th^ % null summed *t* = -3.05. The average Cohen’s *f*^2^ for this cluster was .06, range = [.05 .08]. Additionally, CHP pain intensity was related to the ΔPAF*ΔAP interaction at a right, frontal cluster, 2 channels, average ΔPAF*ΔAP *t* = -2.50, range = [-2.44 -2.55], summed *t* = -4.99, 97.5^th^ % null summed *t* = -2.99, average *f^2^* = .02, range = [.01 .03], which reflected that greater pain was associated with either AP Increases and PAF speeding or AP decreases and PAF slowing. No significant clusters were identified for ΔPAF and results were qualitatively similar when the analyses were rerun using a wider 8-12 Hz PAF calculation range (Supplementary Figure 7).

No channels demonstrated consistent effects of ΔPAF, ΔAP, ΔAP*ΔPAF, or Visit on pain intensity across PHP and CHP.

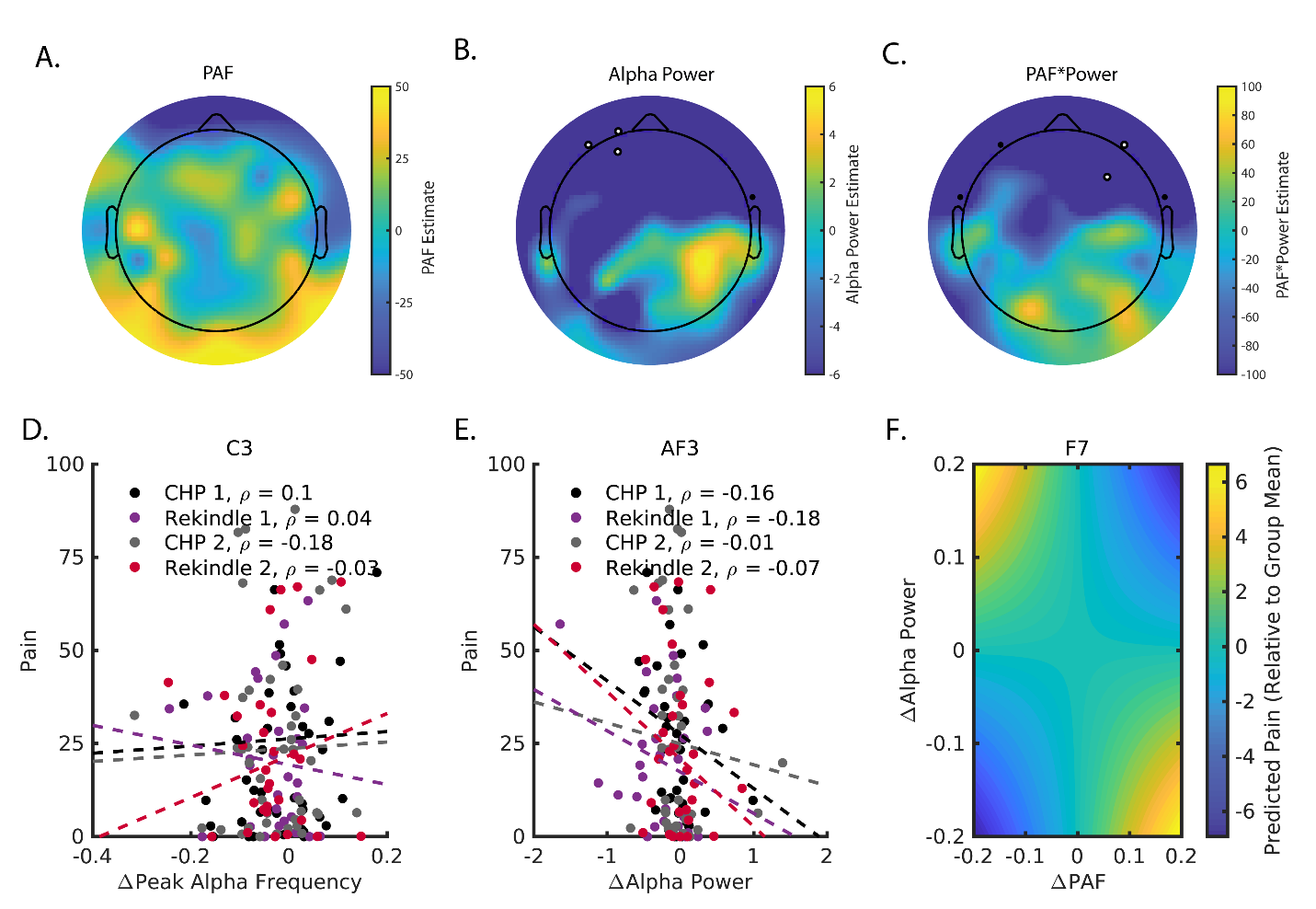

**Supplementary Figure 7.** Alpha Power changes (ΔAlpha Power), but not Peak Alpha Frequency changes (ΔPAF), are scaled by pain intensity during CHP and CHP Rekindle. **A.** No significant channel clusters were identified for ΔPAF. As can be seen in **D.**, there is no reliable relationship between C3 ΔPAF and pain intensity. **B.** ΔAlpha Power is significantly related to pain intensity at a left, frontocentral cluster. As can be seen in **E.**, this effect reflects that increases in power are associated with greater pain. **C.** Pain intensity was related to the ΔPAF*ΔAlpha Power interaction at a right, frontocentral cluster. As shown in **F.**, slowing of PAF with concurrent increases in Alpha Power was associated with greater pain while PAF speeding and power increases were associated with less pain. Data was generated by multiplying hypothetical changes to ΔPAF and ΔAlpha Power with the interaction β obtained from the linear mixed model described in the text. For all topoplots, black dots reflect sensor-level effects surpassing the initial cluster formation threshold (p < .017). Black dots with white centers reflect channels belonging to a significant cluster.

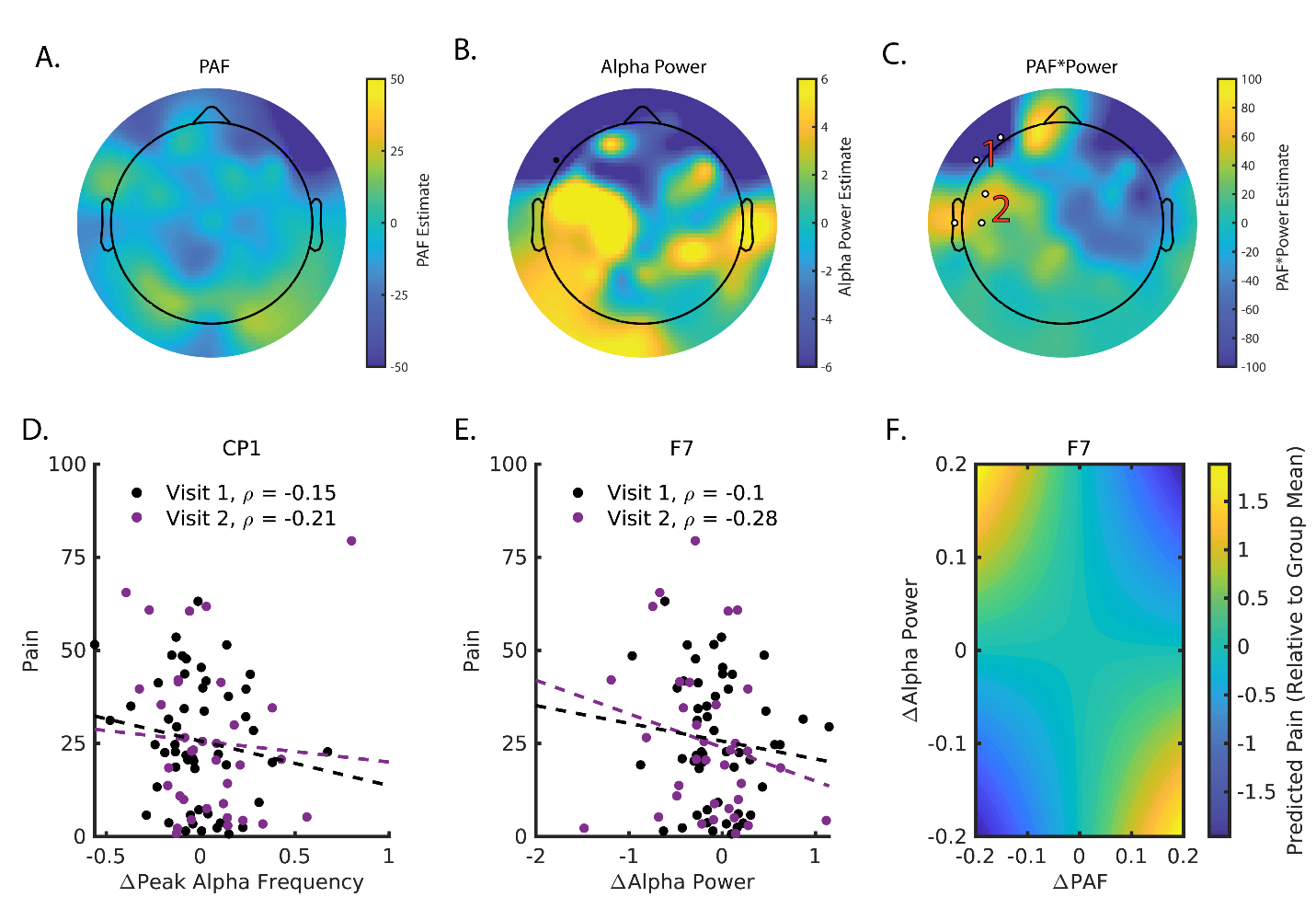
*Prolonged Pain Intensity is not Reliably Scaled by Changes in 8-12 Hz PAF*

**Supplementary Figure 8.** When calculating PAF using the wider 8-12 Hz rang, changes in PAF (ΔPAF) and Alpha Power (ΔAlpha Power) during PHP remain unrelated to pain intensity. No significant clusters were associated with either changes in PAF or Power (**A. & B.**). Scatter plots in **D. & E.** reflect simple correlations between PHP_On_ pain intensity and spectral changes taken from channels surpassing the cluster forming threshold in the original 9-11 Hz PAF analysis. **C.** Pain intensity remains related to the ΔPAF*ΔAlpha Power interaction at a left, frontocentral cluster (1) and the relationship between changes and pain intensity (**F)** is identical to what is seen in Figure 4F. Additionally, we identified a second significant cluster over temporal channels that reflected an inverted relationship to what is seen in Panel F. For all topoplots, black dots reflect sensor-level effects surpassing the initial cluster formation threshold but not belonging to a significant cluster (see Statistics section for specific details on cluster formation for PHP and CHP analyses). White dots outlined with black reflect channels belonging to a significant cluster. Red numbers are used to denote instances where multiple, significant clusters were identified. If no numbers are present within a panel, all outlined channels belong to the same cluster.

Substituting 8-12 Hz PAF estimations in place of 9-11 Hz estimations revealed that PHP pain intensity was again significantly predicted by the ΔPAF x ΔAP at a left, frontal cluster, 2 channels, average ΔPAF x ΔAP *t* = -3.00, range = [-2.79 -3.20], summed *t* = -5.99, 97.5^th^ % null summed *t* = -2.75. Additionally, we identified a novel a left, temporal cluster, 3 channels, average ΔPAF x ΔAP *t* = 2.84, range = [2.53 3.07], summed *t* = 8.51, 97.5^th^ % null summed *t* = 3.18 (Supplementary Figure 8). These effects reflected that at the frontal cluster, greater pain was associated with either AP decreases or PAF speeding or AP increases and PAF slowing while the reverse was true at the temporal cluster. On average, the Cohen’s *f*^2^ associated with the effect of ΔPAF x ΔAP was .09, range = [.08 11], and .09, range = [.07 .14], for the frontal and temporal clusters, respectively. No significant clusters were associated with the ΔPAF or ΔAP.

For CHP pain intensity (Supplementary Figure 9), substituting 8-12 Hz PAF estimations in place of 9-11 Hz estimations failed to reveal any significant clusters for the effects of ΔPAF, ΔAP, or ΔPAF*ΔAP. Despite this, the effect of ΔAP at the originally identified right, frontal cluster was similar but weaker than what we found previously, average ΔAP *t* = -2.16, *t* range = [-1.90 -2.17], summed *t* = -6.50, 97^th^ % summed *t* = -3.05. The effect of ΔPAF*ΔAP at the right, frontal cluster did not, however, mirror our prior results, average ΔAP *t* = .11, *t* range = [-.10 .32], summed *t* = .22, 97^th^ % summed *t* = -3.16.

**Supplementary Figure 9.** When calculating PAF using the wider 8-12 Hz rang, changes in PAF (ΔPAF) and Alpha Power (ΔPower) during CHP remain unrelated to pain intensity. No new significant clusters were associated with ΔPAF, ΔAlpha Power, or ΔPAF*ΔPower (**A - C.**). Scatter plots in **D. & E.** reflect simple correlations between CHP and CHP Rekindle pain intensity and spectral changes taken from channels surpassing the cluster forming threshold in the original 9-11 Hz PAF analysis. **C.** Pain intensity was not related to the ΔPAF*ΔPower interaction at the originally identified right, frontal cluster. **F.** For demonstration purposes the relationship between the interaction and pain intensity at the one channel, F7, surpassing the initial cluster form threshold is shown. For all topoplots, black dots reflect sensor-level effects surpassing the initial cluster formation threshold but not belonging to a significant cluster (see Statistics section for specific details on cluster formation for PHP and CHP analyses). White dots outlined with black reflect channels belonging to a significant cluster. Red numbers are used to denote instances where multiple, significant clusters were identified. If no numbers are present within a panel, all outlined channels belong to the same cluster.

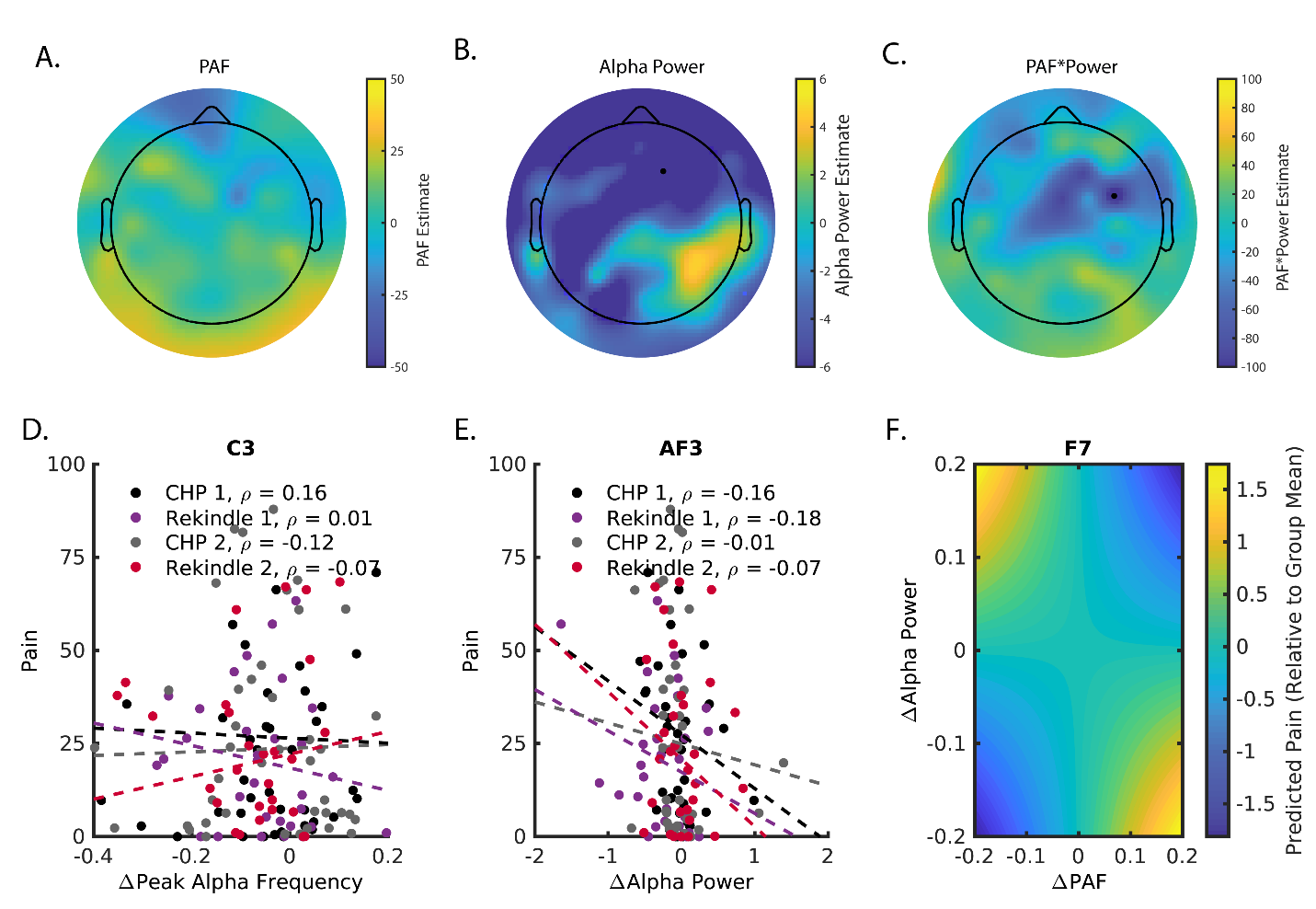

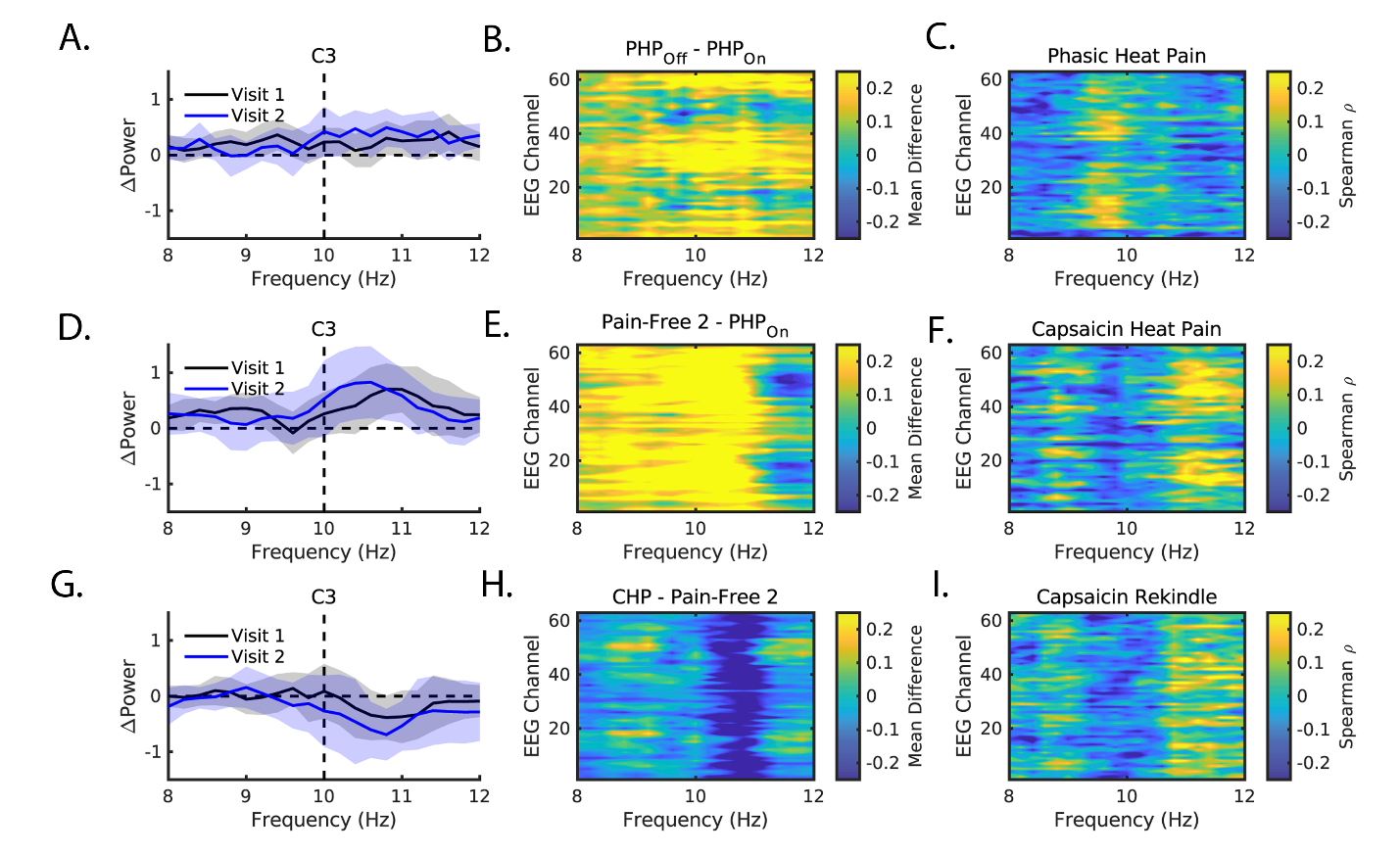

**Supplementary Figure 10. A.** Differences between PHP_Off_ and Pain-Free 1 spectra at Visit 1 (Black) and Visit 2 (Blue). Data is taken from the C3 channel and shaded regions reflect the 95^th^ confidence interval. No clear changes in “fast” or “slow” spectral power are evident. **B.** Same as in A, but with each row representing one of the 63 EEG channels. Increasingly warmer colors indicate greater power increases while increasingly cooler colors indicate greater power decreases. Slight increases in the “fast” alpha range (somewhat stronger in the 10 – 11 Hz range) are seen at nearly all channels. **C.** Correlation between PHP_Off_ power changes (PHP_Off_ – Pain-Free 1) throughout the 8-12 Hz range for each EEG channel. Increasingly warmer colors indicate a greater positive correlation while increasingly cooler colors indicate a greater negative correlation; for ease of visualization, figures represent correlation values averaged across Visits 1 and 2. No clear pattern is evident. **D-F.** Same as in A-C, except with spectral differences between Pain-Free 2 and PHP_On._ Focal increases in “fast” alpha power during Pain-Free 2 occur in a range similar to the one demonstrating power decreases during PHP_On_. **G-I.** Same as in A-C, except with spectral differences between Pain-Free 3 and Pain-Free 2. Decreases in power are once again seen in the “fast” alpha range, indicating that these decreases are not directly tied to the presence of thermode generated heat.
